# Engineered resistance to Zika virus in transgenic *Ae. aegypti* expressing a polycistronic cluster of synthetic miRNAs

**DOI:** 10.1101/344697

**Authors:** Anna Buchman, Stephanie Gamez, Ming Li, Igor Antoshechkin, Shin-Hang Lee, Shin-Wei Wang, Chun-Hong Chen, Melissa J. Klein, Jean-Bernard Duchemin, Prasad N. Paradkar, Omar S. Akbari

## Abstract

Recent Zika virus (ZIKV) outbreaks have highlighted the necessity for development of novel vector control strategies to combat arboviral transmission, including genetic versions of the sterile insect technique, artificial infection with *Wolbachia* to reduce population size and/or vectoring competency, and gene drive based methods. Here, we describe the development of mosquitoes synthetically engineered to impede vector competence to ZIKV. We demonstrate that a polycistronic cluster of engineered microRNAs (miRNAs) targeting ZIKV is expressed and fully processed following a blood meal in *Ae. aegypti*, ensuring the formation of mature synthetic miRNAs in the midgut where ZIKV resides in the early stages of infection. Critically, we demonstrate that engineered *Ae. aegypti* mosquitoes harboring the anti-ZIKV transgene have significantly reduced viral infection, dissemination, and transmission rates of ZIKV. Taken together, these compelling results provide a promising path forward for development of effective genetic-based ZIKV control strategies, which could potentially be extended to curtail other arboviruses.

**One Sentence Summary:** Here we describe the generation of *Ae. aegypti* mosquitoes that are engineered to confer reduced vector competence to Zika virus (ZIKV) and we discuss how such engineering approach can be used to combat the major health burden of ZIKV and potentially other arboviruses in the future.

## Main

Since being introduced into the Americas, ZIKV - a mosquito-borne flavivirus - has spread rapidly, causing hundreds of thousands of cases of ZIKV infection ^1^. Although most cases remain asymptomatic, infection during pregnancy has been associated with severe congenital abnormalities and pregnancy loss, presenting an unprecedented health threat with long-term consequences ^2^. This prompted the World Health Organization to declare ZIKV a Public Health Emergency of International Concern in 2016 ^1,2^. Currently, there are no clinically approved vaccines to prevent ZIKV and no effective treatment options for infected individuals; thus, vector control remains essential in curtailing the ZIKV epidemic. Like dengue virus (DENV) and chikungunya virus (CHIKV), ZIKV is transmitted primarily by *Aedes* mosquitoes, which are expanding their habitable range due to urbanization, climate change, and global trade ^3^. Current methods of vector control, including removal of standing water and use of insecticides, have not been entirely effective in the fight against the spread of *Aedes* mosquitoes ^3^. Therefore, new game-changing innovative vector control strategies, including those utilizing genetically engineered mosquitoes ^4^, are urgently needed to combat the spread of ZIKV and other *Aedes*-vectored diseases worldwide.

Employment of genetically modified (or otherwise altered) insects to manipulate disease-vectoring populations was first proposed decades ago ^5^, and due in part to enabling technological advances, has garnered increased interest in recent years ^4^. In fact, several strategies for genetic-based vector control are currently being utilized in the field. For example, the RIDL (Release of Insects carrying a Dominant Lethal) system ^6^ - a genetic-based Sterile Insect Technique (SIT)-like system - has recently been shown to be effective in reducing wild insect populations ^7^. Open field release trials of these genetically modified mosquitoes have been conducted in several countries, including Cayman Islands, Malaysia, Brazil, and are currently being considered for use in India and the USA ^8–10^. In addition to genetic-based vector control strategies, mosquitoes harboring artificially acquired strains of *Wolbachia* can also be used to either reduce total insect populations ^11^ or to render insect populations less competent vectors of certain viruses, including ZIKV ^12^, and this technique has also been trialed in multiple countries to reduce impact of mosquito-borne diseases ^13–15^. However, given the accumulating evidence that *Wolbachia* can enhance certain flavivirus infections ^16–18^ may lead do reevaluation of this technique. Nevertheless, while current approaches are effective, they require inundative releases of large numbers of insects, which can be laborious and expensive and can impede scalability and worldwide adoption.

Another category of genetic-based vector control involves an engineered gene drive system that can force inheritance in a super-Mendelian fashion, enabling it to increase itself - and any linked “cargo” genes - in frequency with each generation even without conferring fitness advantages to its host ^4,19^. Such a method could be used to disseminate desirable “cargo” genes, such as pathogen resistance, rapidly through wild disease-transmitting populations replacing vector populations with disease-refractory ones ^20^. While significant efforts are currently underway to develop engineered drive systems ^21–24^, others are focused on creation of “cargo” genes that may be spread by the drive systems, and several studies have reported on the successful development of pathogen resistance “cargo” genes in *Ae. aegypti* ^25–27^.

To date, however, no anti-ZIKV refractory “cargo” genes in any mosquito have been developed. To fill this void, here we describe the generation of a synthetically engineered ZIKV resistance transgene comprising a polycistronic cluster ZIKV-targeting synthetic miRNAs. We demonstrate that *Ae. aegypti* mosquitoes harboring this anti-ZIKV transgene express and fully process the ZIKV-targeting synthetic miRNAs in the midgut and consequently have significantly reduced viral infection, dissemination, and transmission rates of ZIKV. Specifically, we demonstrate that mosquitoes homozygous for the anti-ZIKV transgene are fully resistant to ZIKV infection and are unable to transmit the virus. In contrast, we determine that a minority of heterozygotes for the anti-ZIKV transgene can become infected with ZIKV following exposure. However, these heterozygotes become infected at significantly lower rates than wildtypes (WT), and those susceptible to infection have roughly three orders of magnitude lower viral titres in their saliva, suggesting a significantly reduced possibility of viral transmission. This is supported by our finding that heterozygous anti-ZIKV mosquitoes are almost entirely incapable of *in vivo* ZIKV transmission in a sensitive *Stat1-/-* mouse model. Moreover, when compared to *Wolbachia w* Mel positive mosquitoes, previously shown to have reduced ZIKV vectoring competency ^12^, the anti-ZIKV mosquitoes perform siginficantly better in the ZIKV challenge assays. Taken together, these compelling results provide a promising path forward for development of effective ZIKV control - and possibly control of other clinically significant arboviruses - using genetically engineered mosquitoes.

## Results

### Engineering ZIKV-resistant mosquitoes

To generate a “cargo” gene that can confer resistance to ZIKV, we implemented a synthetic miRNA-based approach, since such an approach has been previously demonstrated to generate virus-resistance phenotypes in a number of contexts (e.g., ^28–30^), including mosquitoes ^31^. We engineered a *piggyBac* vector comprising a polycistronic cluster of eight ZIKV-targeting synthetic miRNAs (anti-ZIKV transgene) under control of the *Ae. aegypti* carboxypeptidase promoter ^32^ to drive expression of the synthetic miRNAs in female midguts following a blood meal (Figure 1A). To ensure effective viral suppression and evolutionary stability, we designed each of the eight synthetic miRNAs to target 6/10 conserved protein-coding genes of French Polynesia ZIKV strain H/PF/2013 (GenBank: KJ776791.2) ^33^, including all three structural genes (capsid (C), membrane precursor (prM), envelope (E)), and three non-structural genes (NS1, NS2A, NS5). Each of these genes was targeted by a single synthetic miRNA, except for the RNA-dependent RNA polymerase NS5, which was targeted by three miRNAs due to its importance for the replication of the flaviviral RNA genome (Figure 1B, Supplementary Figure 1). The engineered anti-ZIKV transgene (termed plasmid OA959C) also contained the eye-specific 3xP3 promoter ^34^ driving expression of tdTomato as a transgenesis marker (Figure 1A).

**Figure 1.**
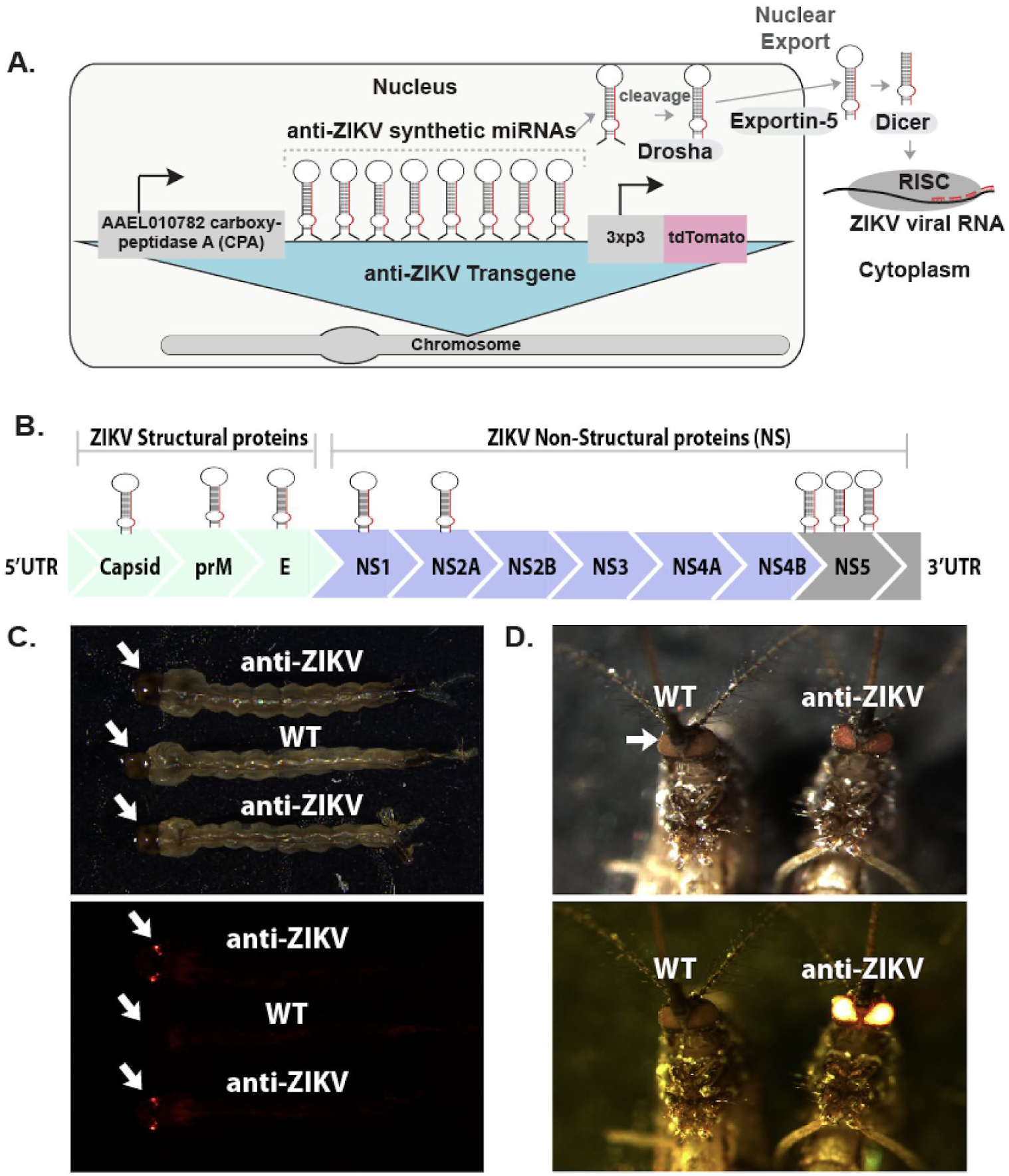
Schematic of anti-ZIKV transgene, ZIKV target sites and phenotype of transgenic mosquitoes. A schematic of the anti-ZIKV transgene used in the study consisting of a carboxypeptidase A (AAEL010782) promoter (CPA) driving expression of a polycistronic cluster of eight synthetic miRNAs engineered to target conserved genes in the ZIKV genome. Following processing using the Drosha/exportin-5/Dicer pathway and loading into the RNA-induced silencing complex (RISC), the miRNAs and their target ZIKV viral RNA interact in the cytoplasm (**A**). A schematic of the ZIKV genome consisting of three structural proteins (Capsid, prM, E) and seven non-structural proteins (NS1, NS2A, NS2B, NS3, NS4A, NS4B and NS5) with relative synthetic miRNA targets indicated by hairpin above (**B**). WT and TZIKV-C larvae (**C**) and adult mosquitoes (**D**) imaged under both transmitted light and a fluorescent dsRED filter.

Following embryonic microinjection, multiple transgenic lines were identified (n > 6), and four independent lines with strong expression of tdTomato fluorescence in the eyes (termed TZIKV-A, B, C, and D) were selected for further characterization (see Figure 1C, 1D for fluorescence in TZIKV-C). To verify the transgene insertion sites, we performed inverse PCR (iPCR) on genomic DNA extracted from transgenic mosquitoes of all four independent strains. iPCR analysis indicated that insertion sites were on chromosome 2 (at approximate position 167,899,561) for line TZIKV-A, on chromosome 3 (at approximate position 402,525,313) for line TZIKV-B, on chromosome 3 (at approximate position 173,647,983) for line TZIKV-C, and on chromosome 1 (at approximate position 228,972,549) for line TZIKV-D when aligned to the AaegL5 assembly (GenBank assembly accession: GCA_002204515.1) ^35^. To avoid any bias due to position effect variegation (PEV) stemming from transgene insertion sites, all four lines were screened for midgut infection status at day 4 post-infection (dpi), and results showed that all 4 lines had significant reduction in midgut infection rate and viral titres compared with WT mosquitoes (Supplementary Figure 2). Given that there was no significant difference in ZIKV reduction in midgut infection between the four lines (TZIKV-A, B, C, and D), the line exhibiting the strongest antiviral phenotype (TZIKV-C) was selected for further comprehensive characterization (Supplementary Figure 2).

### Molecular analysis of synthetic miRNA expression and processing

To confirm expression and processing of the ZIKV-targeting synthetic miRNAs in TZIKV-C, we deep-sequenced small RNA populations from dissected midgut tissues isolated from blood-fed and non-blood-fed female mosquitoes using an Illumina platform. We detected robust expression of the passenger ^36^ and mature miRNA guide strands of 5 out of 8 anti-ZIKV-targeting synthetic miRNAs (miRNAs 1,2,4,6,8) with TPM (transcripts per million) values ranging from 2 to 91, 25.7 on average, indicating that these synthetic miRNAs are efficiently expressed and processed (Supplementary Figure 3, Supplementary Table 1). Importantly, no anti-ZIKV-targeting miRNAs (>1 read) were identified in small RNA populations derived from WT *Ae. aegypti* (Supplementary Table 1). Taken together, these results demonstrate that the anti-ZIKV transgene is stably integrated into the mosquito genome and most of the ZIKV-targeting synthetic miRNAs are properly expressed and processed in an appropriate context (i.e., in midgut) for ZIKV suppression.

### Engineered mosquitoes are refractory to multiple ZIKV strains

To characterize the functional significance of ZIKV-targeting synthetic miRNA expression and processing on vector competence, adult female mosquitoes (WT control and TZIKV-C) were infected with ZIKV (FSS13025, Cambodia 2010 strain, GenBank JN860885) via membrane blood-feeding ^37^. For these experiments we used the Cambodia ZIKV strain, which is from the Asian ZIKV lineage and in close phylogenetic proximity to the French Polynesia ZIKV strain, against which the miRNAs were designed ^38^. Importantly, seven out of eight of the ZIKV-targeting synthetic miRNA target sites are 100% conserved between the Cambodia ZIKV strain and the French Polynesia strain, allowing either strain to be used for the ZIKV challenges (Supplementary Figure 1). At 4 days post-infection (dpi), midguts from blood-fed mosquitoes were dissected and ZIKV titres were measured using real-time RT-qPCR. Results from 3 biological replicates revealed that none of the TZIKV-C mosquitoes homozygous for the transgene (n=32) were positive for ZIKV infection in the midguts. ZIKV infection was detected in 87.5% (28/32) of the TZIKV-C mosquitoes that were heterozygous for the transgene; however, these mosquitoes had significantly (p<0.001) lower viral RNA levels (~2 logs) than WT (Figure 2A, Table 1).

**Figure 2.**
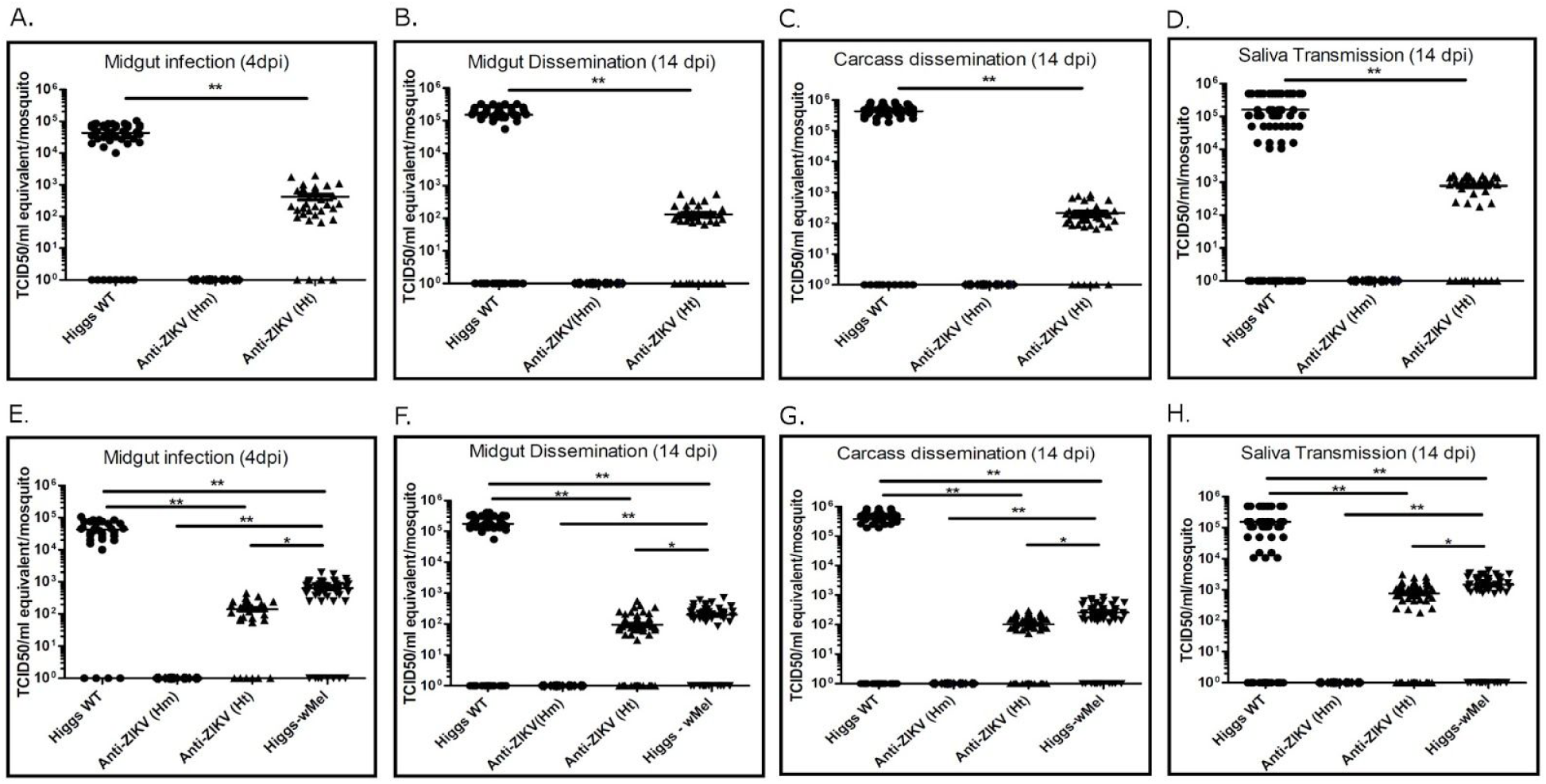
ZIKV titres in WT, TZIKV-C and *Wolbachia* infected mosquitoes challenged with either a Cambodian or a Puerto Rican ZIKV strain. ZIKV virus titres in wildtype (WT), anti-ZIKV homozygous (Hm) and heterozygous (Ht) transgenic mosquitoes, and Wolbachia-infected WT (Higgs-*w*Mel) following a blood meal infected with a Cambodian (FSS13025, **A**-**D**) or Puerto Rican (PRVABC59, **E**-**H**) strain of ZIKV are shown. Virus titres (TCID_50_/ml equivalent) from mosquito midgut (day 4 (**A**, **E**) and day 14 (**B**, **F**) post infection, dpi) and carcass (14 dpi (**C**, **G**)) of WT and transgenic mosquitoes were determined using real-time RT-qPCR and calculated using previously published methods. Virus titres in the saliva collected from WT and transgenic mosquitoes 14 dpi were determined using TCID_50_ on Vero cells and plotted (**D**, **H**). Circles represent WT mosquitoes; diamonds represent anti-ZIKV Hm transgenic mosquitoes; triangles represent TZIKV-C Ht mosquitoes; upside down triangles represent Higgs-*w*Mel mosquitoes. Horizontal bars represent the mean virus titre. *represents p<0.05 and **represents p<0.001. For each experiment, data from three replicates are pooled.

**Table 1.**
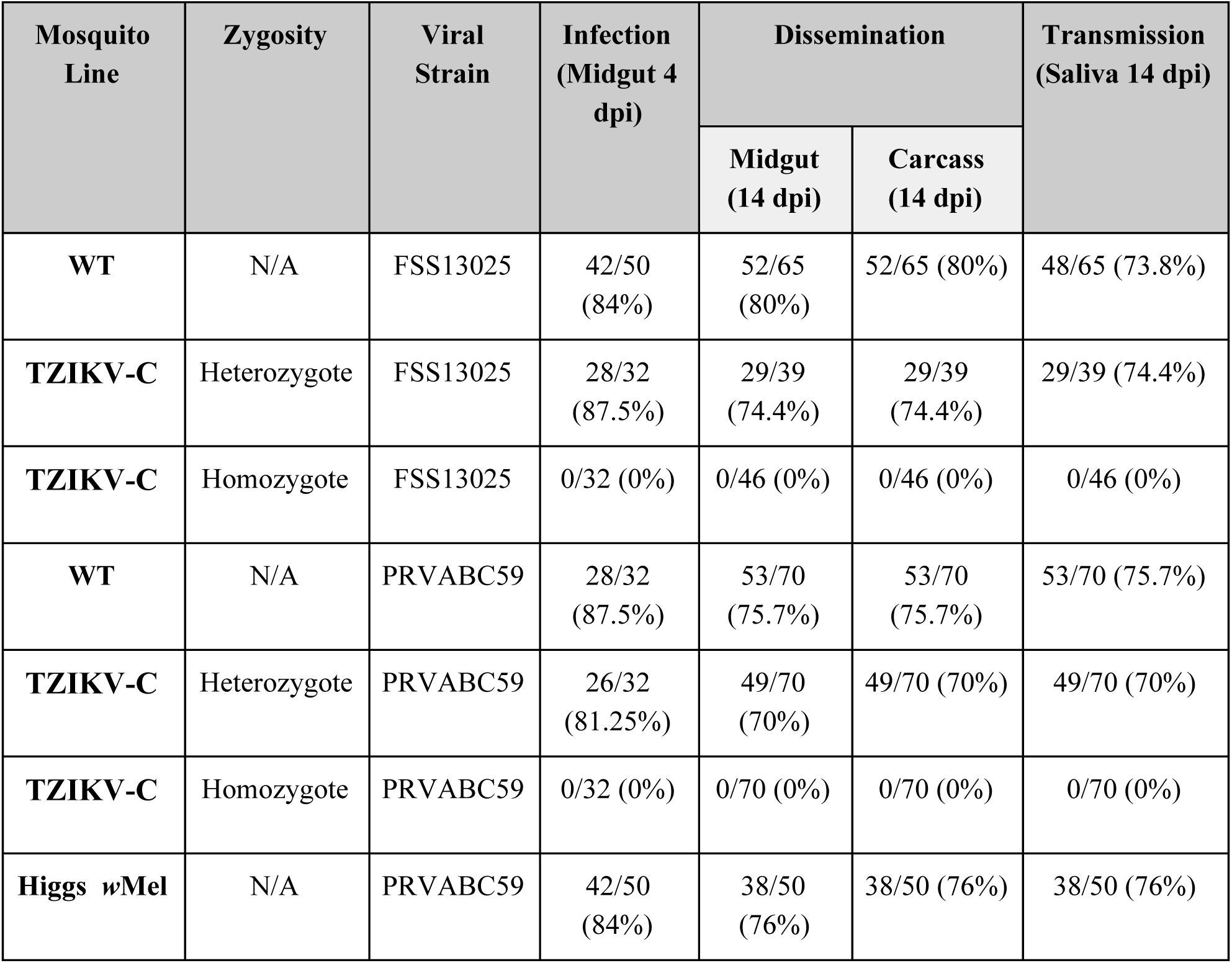
Anti-ZIKV transgene effect on ZIKV infection, dissemination, and transmission rates. ZIKV infection rates were quantified in the midgut at 4 days post infection (dpi). Dissemination rates were quantified in both the midgut and carcass at 14 dpi. Transmission rates were calculated by measuring prevalence of ZIKV in the saliva at 14 dpi. For each experiment, data from three replicates is pooled.

To assay for viral dissemination, total RNA was collected from whole TZIKV-C mosquito carcasses and dissected midguts from both homozygous and heterozygous transgenic mosquitoes at 14 dpi. The results from three biological replicates indicated that none of the homozygous TZIKV-C mosquitoes (n=46) were positive for viral replication (dissemination) in either the midgut or the carcass. ZIKV prevalence was detected in 74.4% (29/39) of heterozygous TZIKV-C mosquitoes in both the carcass and midgut; however, they had significantly (p<0.001) lower levels of viral RNA (~3 logs) compared to WT (Figure 2B-C, Table 1). Finally, to determine viral transmission, saliva from individual mosquitoes was collected at 14 dpi and ZIKV titres were measured using TCID_50_ assay. No ZIKV was detected in the saliva of homozygous TZIKV-C mosquitoes (n=46). Prevalence of ZIKV in the saliva was detected in 74.4% (29/39) of heterozygous TZIKV-C mosquitoes; however, here again the ZIKV titres were significantly (p<0.001) lower (~3 logs) as compared to WT (Figure 2D, Table 1).

To determine whether the synthetic miRNAs are broadly inhibitory for ZIKV, vector competence of transgenic TZIKV-C mosquitoes was also assessed using a second contemporary ZIKV strain (PRVABC59, isolated from US traveller to Puerto Rico in 2015, GenBank KU501215). For this strain, seven out of eight ZIKV-targeting synthetic miRNA target sites (although not the same seven as for the Cambodia strain) are 100% conserved with the French Polynesia strain against which the miRNAs were designed (Supplementary Figure 1). Tests for infection, dissemination, and transmission were carried out as above, and the results were comparable to those obtained with the Cambodia strain. Briefly, at 4 dpi, none of the TZIKV-C mosquitoes homozygous for the transgene (n=32) were positive for ZIKV infection in their midguts, and while ZIKV infection was detected in 81.25% (26/32) of the TZIKV-C mosquitoes that were heterozygous for the transgene, these had significantly (p<0.001) lower viral RNA levels (~2 logs) than WT (Figure 2E, Table 1). TZIKV-C mosquito carcasses and dissected midguts at 14 dpi showed that none of the homozygous TZIKV-C mosquitoes (n=70) were positive for viral replication in either the midgut or the carcass by real-time RT-qPCR, while 70% (49/70) of heterozygous mosquitoes had ZIKV in both the carcass and midgut, albeit with significantly (p<0.001) lower levels of viral RNA (~3 logs) than WT (Figure 2F-G, Table 1). Finally, ZIKV titre measurements on saliva from individual mosquitoes at 14 dpi demonstrated that no ZIKV was present in homozygous TZIKV-C mosquitoes (n=70) indicating that they would be unable to transmit the virus. Prevalence of ZIKV in saliva was detected in 70% (49/70) of TZIKV-C heterozygous mosquitoes; however, here again the ZIKV titres were significantly (p<0.001) lower (~3 logs) as compared with WT (Figure 2H, Table 1).

**Figure 3.**
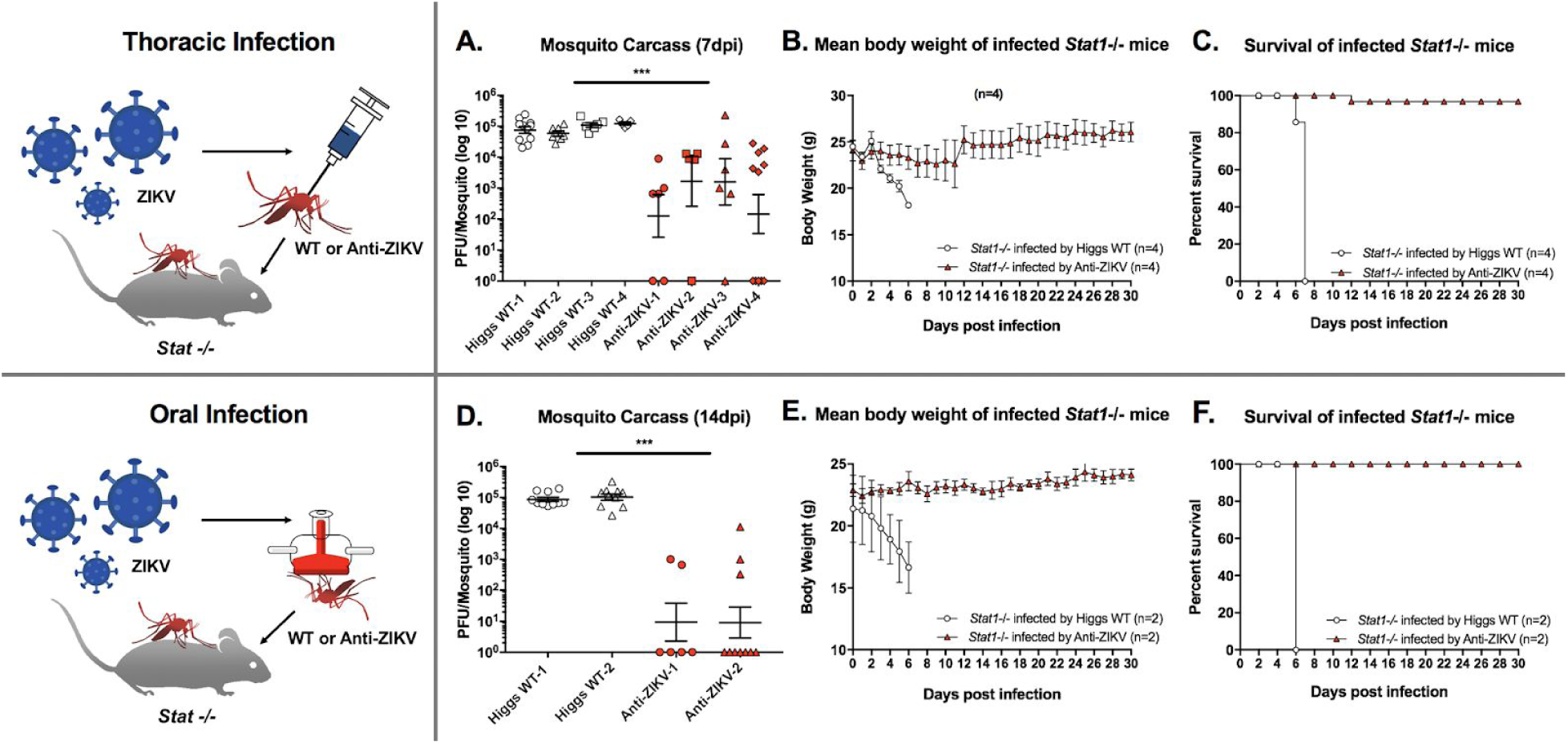
Effect of anti-ZIKV transgene on ZIKV transmission in a mouse model. Wildtype (WT) and heterozygous TZIKV-C mosquitoes were infected with ZIKV strain PRVABC59 thoracically (**A-C**) or orally (**D-F**), and assayed for their ability to transmit ZIKV to immunocompromised *Stat -/-* mice. Viral titres in carcasses of mosquitoes infected thoracically (measured at 7 dpi, **A**) and orally (measured at 14 dpi, **D**) were determined by plaque assay in Vero cells and plotted. Mean body weight (**B, E**) and survival (**C**, **F**) of *Stat -/-* mice following ZIKV infection by thoracially (**B**, **C**) or orally (**E, F**) infected WT and TZIKV-C mosquitoes were measured and plotted. For all plots **(A-F),** white shapes represent results from WT mosquitoes; red shapes represent TZIKV-C mosquitoes. For viral titre plots, horizontal bars represent the mean virus titre, and vertical bars represent SEM. For mean body weight plots, vertical bars represent SEM. ***represents p<0.0001.

### Engineered mosquitoes outperform *Wolbachia*

We next compared the inhibitory effect of our synthetic miRNAs to ZIKV inhibition previously shown with *Wolbachia ^12,39–41^*. Vector competence results revealed that midguts from mosquitoes (Higgs strain) infected with *Wolbachia* (*w*Mel strain; n=50) had significantly (p<0.001) reduced ZIKV (Puerto Rican strain) viral titres (~2 logs) at 4 dpi compared with WT (n=32; Figure 2E, Table 1). Similarly, viral dissemination at 14 dpi was also reduced (p<0.001) in *w*Mel mosquitoes (~3 logs, n=50 Figure 2F-G, Table 1), and ZIKV titres in mosquito saliva at 14 dpi were significantly (p<0.01) lower (~2 logs) in *w*Mel mosquitoes than in WT (Figure 2H, Table 1). Importantly, comparison to the TZIKV-C mosquitoes revealed that the TZIKV-C mosquitoes are significantly (p<0.001) more effective as homozygotes, and modestly more effective as heterozygotes (p<0.05), at blocking ZIKV infection compared with *Wolbachia*-infected mosquitoes.

### Anti-ZIKV transgene inhibits ZIKV transmission in a mouse model

To further characterize ZIKV inhibition by the anti-ZIKV transgene, we also conducted limited tests of *in vivo* transmission capacity on heterozygous TZIKV-C mosquitoes. Specifically, we utilized a very sensitive STAT knockout (*Stat1* -/-) mouse model, in which challenge with ZIKV (either intraperitoneally or via feeding by infected mosquito) rapidly causes systemic infections presenting high virema, resulting in significant weight loss, brain infections, and mortality ^42^. Firstly, we infected adult female mosquitoes (WT and TZIKV-C) with Puerto Rican ZIKV strain (PRVABC59) via thoracic injection, which bypasses the midgut barrier resulting in a significant viral titre in mosquitoes ^42^. At 7 dpi, TZIKV-C (n=28) and WT (n=29) mosquitoes were separately pooled into four groups (with 6-12 individuals per group; Figure 3A) and each group was allowed to blood feed on a *Stat1* -/- mouse, after which mouse weight and survival were measured daily. All mice fed on by infected WT mosquitoes (n=4) became viremic and died prior to 8 dpi with significant weight loss prior to death (p<0.05; Figure 3B, C). Conversely, out of the four mice fed on by TZIKV-C mosquitoes, only one showed mortality (albeit at a later date - 12 dpi), and no significant weight loss was observed compared to the control group (p<0.0001; Figure 3B, C). Measurement of ZIKV titres in the individual mosquitoes utilized for this assay demonstrated that nearly all TZIKV-C mosquitoes had significantly reduced viral titers (~2 log) at 7 dpi compared to WT (p<0.0001; Figure 3A).

To better simulate how mosquitoes naturally obtain pathogens (i.e., from blood feeding), we also performed the above assay with mosquitoes that were infected with ZIKV (strain PRVABC59) via oral membrane blood feeding, and obtained similar results. Specifically, at 14 dpi via oral membrane blood feeding, TZIKV-C (n=16) and WT (n=20) mosquitoes were pooled into groups of 6-10 and allowed to feed on, and transmit the virus to, *Stat1*-/- mice (n=2 for each transgenic and WT; Figure 3D). The mice fed on by infected WT mosquitoes experienced significant weight loss and mortality prior to 8 dpi (p<0.0001; Figure 3E, F). Conversely, mice fed on by TZIKV-C mosquitoes showed no significant change in weight and no infection-associated mortality (p<0.0001, Figure 3E, F). Viral titre assays on these mosquitoes (at 14 dpi) indicated that ZIKV infection rate was dramatically reduced in TZIKV-C mosquitoes (39% of mosquitoes infected compared to 93% of WT; Figure 3D), and that viral titres of the TZIKV-C mosquitoes that were infected were significantly lower (~2 log) than that of WT (p<0.0001, Figure 3D). Altogether, these results demonstrate that the anti-ZIKV transgene confers robust refractoriness to ZIKV infection and transmission, and that even mosquitoes heterozygous for the transgene are unlikely to be able to transmit the virus.

### Impact of anti-ZIKV transgene on mosquito fitness

Finally, to determine whether the anti-ZIKV transgene had any significant fitness effects on transgenic mosquitoes, we assessed several fitness parameters including larval to pupal development time, male female fecundity and fertility, male mating success, adult wing length, and longevity (Table 2). No significant differences between WT and TZIKV-C mosquitoes were observed when examining male mating success, fecundity, and fertility (all *p* values > 0.9); female fecundity (*p*>0.05); and male and female wing length (*p*>0.05). Conversely, we observed a significant difference (*p*<0.01) in hatching rates of eggs laid by WT versus TZIKV-C females (with the latter having lower hatching rates), and a significant difference between larval to pupal development time (*p*< 0.01), with TZIKV-C individuals developing faster. When assessing adult mosquito survivorship, no significant differences were observed between WT and TZIKV-C males (*p*>0.05; Table 2, Supplementary Figure 4), while WT females survived slightly longer than TZIKV-C females (*p*<0.0001; Table 2, Supplementary Figure 4). Furthermore, there was no significant difference in survival at 14 dpi between WT and TZIKV-C mosquitoes infected with the Cambodia ZIKV strain (*p*>0.05; Table 2, Supplementary Table 2), and similarly no significant difference in survival between WT, *w*Mel infected, and TZIKV-C mosquitoes infected with the Puerto Rico ZIKV strain (*p*>0.05; Table 2, Supplementary Table 2). Based on the above observations, it does not appear that the anti-ZIKV transgene imposes any obvious major fitness costs to its carriers.

**Table 2.**
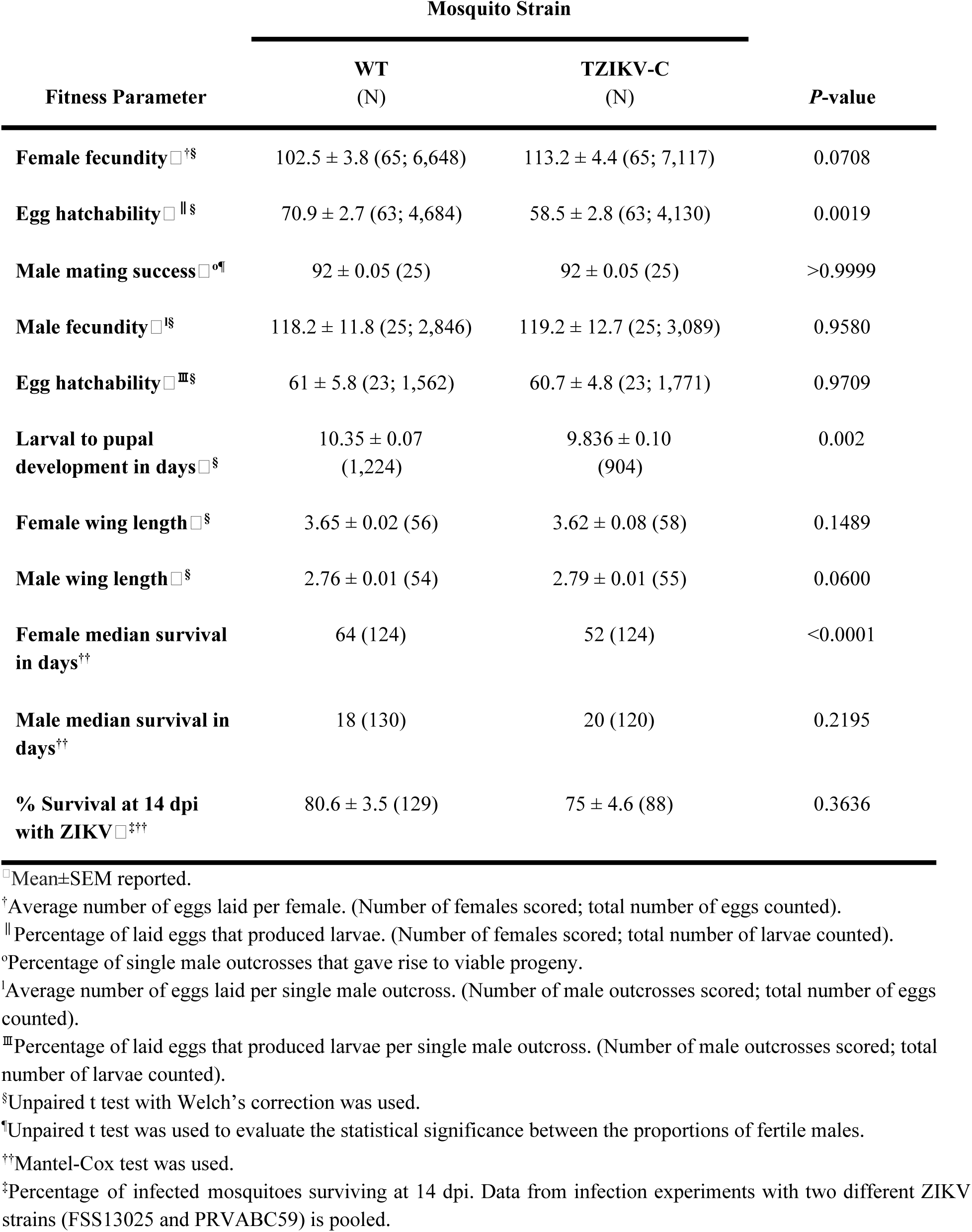
Fitness evaluation of WT and TZIKV-C mosquitoes. Comparisons of several fitness parameters (leftmost column) between WT (second column from left) and TZIKV-C mosquitoes (third column from left) suggest that there are few significant differences (rightmost column) between the two groups, indicating that the ani-ZIKV transgene does not have a major impact on mosquito fitness.

| Fitness Parameter | Mosquito Strain |  | *P*-value |
| --- | --- | --- | --- |
|  | WT<br>(N) | TZIKV-C<br>(N) |  |
| **Female fecundity**□<sup>†§</sup> | 102.5 ± 3.8 (65; 6,648) | 113.2 ± 4.4 (65; 7,117) | 0.0708 |
| **Egg hatchability**□<sup>‡§</sup> | 70.9 ± 2.7 (63; 4,684) | 58.5 ± 2.8 (63; 4,130) | 0.0019 |
| **Male mating success**□<sup>°¶</sup> | 92 ± 0.05 (25) | 92 ± 0.05 (25) | >0.9999 |
| **Male fecundity**□<sup>‡§</sup> | 118.2 ± 11.8 (25; 2,846) | 119.2 ± 12.7 (25; 3,089) | 0.9580 |
| **Egg hatchability**□<sup>‡‡§</sup> | 61 ± 5.8 (23; 1,562) | 60.7 ± 4.8 (23; 1,771) | 0.9709 |
| **Larval to pupal development in days**□<sup>§</sup> | 10.35 ± 0.07<br>(1,224) | 9.836 ± 0.10<br>(904) | 0.002 |
| **Female wing length**□<sup>§</sup> | 3.65 ± 0.02 (56) | 3.62 ± 0.08 (58) | 0.1489 |
| **Male wing length**□<sup>§</sup> | 2.76 ± 0.01 (54) | 2.79 ± 0.01 (55) | 0.0600 |
| **Female median survival in days**<sup>††</sup> | 64 (124) | 52 (124) | <0.0001 |
| **Male median survival in days**<sup>††</sup> | 18 (130) | 20 (120) | 0.2195 |
| **% Survival at 14 dpi with ZIKV**□<sup>‡††</sup> | 80.6 ± 3.5 (129) | 75 ± 4.6 (88) | 0.3636 |

□ Mean±SEM reported.
<sup>†</sup> Average number of eggs laid per female. (Number of females scored; total number of eggs counted).
<sup>‡</sup> Percentage of laid eggs that produced larvae. (Number of females scored; total number of larvae counted).
<sup>°</sup> Percentage of single male outcrosses that gave rise to viable progeny.
<sup>‡</sup> Average number of eggs laid per single male outcross. (Number of male outcrosses scored; total number of eggs counted).
<sup>‡‡</sup> Percentage of laid eggs that produced larvae per single male outcross. (Number of male outcrosses scored; total number of larvae counted).
<sup>§</sup> Unpaired t test with Welch's correction was used.
<sup>¶</sup> Unpaired t test was used to evaluate the statistical significance between the proportions of fertile males.
<sup>††</sup>Mantel-Cox test was used.

## Discussion

Taken together, our results demonstrate that targeting conserved genes in the ZIKV genome by expressing an engineered polycistronic cluster of synthetic microRNAs confers homozygous mosquitoes complete refractoriness to multiple strains of ZIKV infection, dissemination, and transmission. Although incomplete, heterozygous mosquitoes also display partial refractoriness to ZIKV infection, dissemination, and transmission, with significant reduction of viral titres in the saliva (>2 logs compared to WT). This significant reduction of ZIKV is greater than the viral inhibition effect of *Wolbachia*, and may be enough to ensure these heterozygous mosquitoes are unable to transmit ZIKV to a susceptible host in the wild. Indeed, this latter point is supported by our finding that heterozygotes were largely unable to transmit ZIKV to immunocompromised (*Stat1 -/-*) mice after infection via thoracic injection, and completely unable to transmit ZIKV after infection via membrane feeding. As intrathoracic injection generates unnaturally high infection levels by bypassing the mosquito midgut and lumen barriers ^42^, it is perhaps unsurprising that one mouse (possibly fed on by the single transgenic mosquito with unusual WT-like viral titre; Figure 3A) experienced ZIKV-associated mortality. However, the observation that most of the heterozygous anti-ZIKV mosquitoes infected via thoracic injection - and all of the mosquitoes infected via membrane feeding - were unable to transmit ZIKV to a susceptible mouse model, strongly suggests that even heterozygotes are unlikely to be capable of ZIKV transmission in the wild.

Previously in *Ae. aegypti*, resistance to DENV has been engineered by transgenic activation of antiviral pathways ^25^, transgene-based RNAi in either the midgut ^26,43^ or salivary glands ^27^, and antiviral hammerhead enzymes ^44^, and expression of synthetic miRNAs has also been demonstrated to induce partial resistance to DENV-3 and CHIKV^31^. However, similar approaches have not been successfully demonstrated for ZIKV; and, the currently described system is potentially especially advantageous, since targeting 6/10 conserved protein-coding genes from the ZIKV genome with 8 separate synthetic miRNAs may reduce the possibility of escapee mutants and thus ensure evolutionary stability. That said, it remains uncertain how many synthetic miRNAs are necessary to ensure evolutionary stability in a wild population. Moreover, in our small RNA sequencing efforts we only detected expression/processing from 5 out of the 8 synthetic miRNAs indicating that perhaps the RISC machinery is overloaded, or perhaps the CPA promoter is not strong enough to ensure robust expression from all 8 synthetic miRNAs, or possibly some synthetic miRNAs are unstable and get quickly degraded after processing. The latter hypothesis is supported by the fact that robust expression was only detected from synthetic miRNAs 1,2,4,6,8, while 3,5,7 were undetected and given that these are arranged numerically in a linear array (i.e. 1-8) miRNAs 3,5,7 must have been expressed/processed in order for 4,6,8 to be expressed/processed/detected. Therefore, future efforts should be focused on addressing these above open questions.

Altogether, this strategy may provide a suitable “cargo” gene for practical use with a gene drive system to reduce/eliminate vector competence of mosquito populations. For example, previous reports have shown that Cas9-mediated homing based gene drive can be used for population modification of the malaria vector mosquito, *Anopheles stephensi ^21^*, and it should be possible to develop similar systems in *Ae. aegypti*. Given that homing based drive systems quickly convert heterozygotes into homozygotes ^4^, linking the anti-ZIKV transgene such as the one described here to such a system could quickly convert an entire mosquito population into anti-ZIKV homozygotes that would be 100% resistant to ZIKV transmission.

Recent ZIKV outbreaks have shown that vector control remains an essential part of reducing the health burden of emerging arboviruses. Although the aim of this study was to illustrate feasibility of producing ZIKV-refractory mosquitoes, similar genetic engineering strategies could be used to develop (or improve on ^31^) single transgenes that render mosquitoes completely resistant to multiple arboviruses like DENV and CHIKV. Given the increasing incidence of these viral infections worldwide, such transgenes (coupled with gene drive systems) can provide an effective, sustainable, and comprehensive strategy for reducing the impact of arboviral mosquito-borne diseases.

## Acknowledgements

This work was supported in part by an NIH-K22 Career Transition award (5K22AI113060), an NIH Exploratory/Developmental Research Grant Award (1R21AI123937) awarded to O.S.A and CSIRO internal funding to P.N.P. We thank Prof. Robert Tesh and Dr. Nikos Vasilakis (UTMB, Texas) for providing the ZIKV FSS13025 and PRVABC59 strains and Lee Trinidad for preparing viral stocks. We thank Christian Bowman for helping with the inverse PCR. We also thank Prof. Scott O’Neill (Institute of Vector Borne Diseases, Monash University, Australia) and World Mosquito Program for providing *Wolbachia* infected mosquito eggs. Finally, we thank Dr. Guann-Yi Yu (NTU, Taiwan) for providing the *Stat -/-* mice used in this study.

## Author Contributions

A.B., P.N.P., and O.S.A conceived and designed experiments. A.B., S.G, M.L. performed all molecular and genetic experiments. I.A performed sequencing and bioinformatic analysis. M.J.K., J.B.D and P.N.P. performed and analyzed data for ZIKV mosquito challenge assays. S.H.L., S.W.W., and C.H.C performed and analyzed data for *in vivo* mouse model ZIKV mosquito challenge assays. All authors contributed to the writing and approved the final manuscript.

## Materials & Correspondence

Correspondence to: &

## Methods

### Synthetic anti-ZIKV miRNAs design and construction

The *Drosophila melanogaster* miR6.1 stem-loop, which has been previously validated in *D. melanogaster* ^23^, was modified to target eight unique sites in the ZIKV polyprotein region as previously described ^24^. The eight target sites corresponded to regions of capsid (C), membrane precursor (prM), and envelope (E) structural genes, RNA-directed RNA polymerase NS5 (which contained three target sites), and non-structural proteins NS1 and NS2A, of ZIKV strain H/PF/2013 (GenBank: KJ776791.2) ^33^. These sites were highly conserved in ZIKV strain FSS13025 (Cambodia 2010, Genbank KU955593)^37^ and in ZIKV strain PRVABC59 (isolated from US traveller to Puerto Rico in 2015, GenBank KU501215) (Supplementary Figure 1). To generate miR6.1 stem-loop backbones that create mature synthetic miRNAs complementary to each of these target sites, pairs of primers were annealed and products were utilized for two subsequent rounds of PCR and cloned into the pFusA backbone (from the Golden Gate TALEN and TAL Effector Kit 2.0, Addgene #1000000024) in sets of four using Golden Gate assembly ^45^ to generate plasmids OA959A and OA959B. Assembled miRNAs were then digested with either PmeI/BglII (vector OA959A) or with BamHI/PacI (vector OA959B) and were subcloned into a PacI/PmeI-digested final vector OA959C (the anti-ZIKV transgene). The ZIKV target sequences and sequences of primers used in the miRNA cloning are listed in Supplementary Table 3.

### Plasmid Assembly

To generate vector OA959C (the anti-ZIKV transgene), several components were cloned into the *piggyBac* plasmid pBac[3xP3-DsRed] ^46^ using Gibson assembly/EA cloning ^47^. First, a *Drosophila* codon Ref46optimized tdTomato marker was amplified with primers 959C.10A and 959C.10B from a gene synthesized vector (GenScript, Piscataway, NJ) and cloned into a XhoI/FseI digested pBac[3xP3-DsRed] backbone using EA cloning. The resulting plasmid was digested with AscI, and the following components were cloned in via EA cloning: the predicted *Aedes aegypti* carboxypeptidase promoter ^32^ amplified from *Ae. aegypti* genomic DNA using primers 959C.11A and 959C.11B, a GFP sequence amplified from vector pMos[3xP3-eGFP] ^48^ with primers 959C.12A and 959C.12B, and a 677 bp p10 3’ untranslated region (UTR) amplified with primers 959C. 13A and 959C.13B from vector pJFRC81-10XUAS-IVS-Syn21-GFP-p10 (Addgene plasmid #36432). Assembled miRNA fourmers were then subcloned into final plasmid OA959C using PacI and PmeI using traditional cloning. All primer sequences are listed in Supplementary Table 4. Complete annotated plasmid sequence and DNA is available via Addgene (plasmid #104968).

### Generation of Transgenic Mosquitoes

Germline transformations were carried out largely as described ^49^. Briefly, 0-1 hr old Higgs and Liverpool strain *Ae. aegypti* pre-blastoderm embryos were injected with a mixture of vector OA959C (200 ng/ul) and a source of *piggyBac* transposase (200 ng/ul) ^48^; the injected embryos were hatched in deoxygenated H_2_O. A total of 52 surviving Higgs adult males and 64 surviving Higgs adult females, and 61 surviving adult Liverpool males and 75 surviving adult Liverpool females, respectively, were recovered after the injection. Higgs adults were assigned to 35 pools and Liverpool adults were assigned to 39 pools, and outcrossed to Higgs or Liverpool adults, respectively, of the opposite sex in cages. Larvae were fed ground fish food (TetraMin Tropical Flakes, Tetra Werke, Melle, Germany) and adults wRef49ere fed with 0.3M aqueous sucrose. Adult females were blood fed three to five days after eclosion using anesthetized mice. All animals were handled in accordance with the guide for the care and use of laboratory animals as recommended by the National Institutes of Health and supervised by the local Institutional Animal Care and Use Committee (IACUC). A total of 8,189 Higgs and 10,949 Liverpool G_1_s were screened. Larvae with positive fluorescent signals (3xp3-tdTomato) were selected under the fluorescent stereomicroscope (Leica M165FC) and were crossed to establish stable transgenic lines. Four independent lines (termed TZIKV-A, B, C, and D) with the strongest fluorescence expression patterns were selected for further characterization. To determine whether these lines represented single chromosomal insertions, we backcrossed single individuals from each of the lines for four generations to wild-type stock, and measured the Mendelian transmission ratios in each generation; in all cases, we observed a 50% transmission ratio, indicating insertion into single chromosomes. For one of the four lines (TZIKV-C), transgenic mosquitoes were inbred for at least 12 generations to generate a homozygous stock. Mosquito husbandry was performed under standard conditions as previously described ^50^.

### Characterization of Transgene Genomic Insertion Sites

To characterize the insertion site of vector OA959C in transgenic mosquitoes, we adapted a previously described inverse polymerase chain reaction (iPCR) protocol ^51^ as follows. Genomic DNA (gDNA) was extracted from 10 transgenic *Ae. aegypti* fourth instar larvae using the DNAeasy Blood & Tissue Kit (Qiagen #69504) per the manufacturer’s protocol. Two separate restriction digests were performed on diluted gDNA to characterize the 5’ and 3’ ends of the insertion using Sau3AI (5’ reaction) or HinP1I (3’ reaction) restriction enzymes. A ligation step using NEB T4 DNA Ligase (NEB #M0202S) was performed on the restriction digest products to circularize digested gDNA fragments, and two subsequent rounds of PCR were carried out per ligation using corresponding *piggyBac* primers listed in Supplementary Table 5. Final PCR products were cleaned up using the MinElute PCR Purification Kit (Qiagen #28004) in accordance with the manufacturer’s protocol, and sequenced via Sanger sequencing (Source BioScience, Nottingham, UK). To confirm transgene insertion locus and orientation via PCR, primers were designed based on iPCR mapped genomic regions and used in tandem with *piggyBac* primers based on their location as listed in Supplementary Table 5. Sequencing data was then blasted to the AaegL5.0 reference genome (NCBI). An alignment of the sequencing data was carried out with SeqManPro (DNASTAR, Madison, WI) to determine orientation of the transgene insertion site. Analysis of the sequencing data indicated that the insertion sites were on chromosome 2 (at approximate position 167,899,561) for line TZIKV-A, on chromosome 3 (at approximate position 402,525,313) for line TZIKV-B, on chromosome 3 (at approximate position 173,647,938) for line TZIKV-C, and on chromosome 1 (at approximate position 228,972,549) for line TZIKV-D. These insertion locations were also confirmed by PCR and sequencing performed on genomic DNA from the transgenic mosquitoes.

### Small RNA Extraction, Isolation, Sequencing, and Bioinformatics

Total RNA was extracted from midguts of 30 ZIKV-C transgenic and WT (Higgs strain) non-blood-fed adult females as well as midguts of 30 ZIKV-C transgenic and WT (Higgs strain) adult females 24 hours post blood-feeding using the Ambion mirVana mRNA Isolation Kit (ThermoFisher Scientific #AM1560). Following extraction, RNA was treated with Ambion Turbo DNase (ThermoFisher Scientific #AM2238). The quality of RNA was assessed using RNA 6000 Pico Kit for Bioanalyzer (Agilent Technologies #5067-1513) and a NanoDrop 1000 UV–vis spectrophotometer (NanoDrop Technologies/Thermo Scientific, Wilmington, DE). Small RNA was then extracted and prepared for sequencing with QIAseq miRNA Library Kit (Qiagen #331502). Libraries were quantified with Qubit dsDNA HS Kit (ThermoFisher Scientific #Q32854) and High Sensitivity DNA Kit for Bioanalyzer (Agilent Technologies #5067- 4626) and sequenced on Illumina HiSeq2500 in single read mode with the read length of 75 nt following manufacturer’s instructions. After adapter trimming and UMI extraction, reads were aligned to mature *Ae. aegypti* miRNAs downloaded from miRBase (release 22, ^52^) and to each synthetic miRNA’s passenger, loop, and guide sequences using bowtie2 in ‘very-sensitive-local’ mode. Custom Perl scripts were used to quantify the number of reads that mapped to each target. Correlation coefficients of TPM values between WT and transgenic animals were calculated in R^53^. Differential expression analysis was performed with R package DESeq2 using two factor design (design= ~ feeding + genotype). TPM values, MA, and volcano plots were generated with R package ggplot2 (Supplementary Figure 3). Quantification data are shown in Supplementary Table 1. All sequencing data can be accessed at NCBI SRA (accsession ID: SRP150144; BioProject ID: PRJNA475410).

### ZIKV Infection of Mosquitoes, Virus Determination and Longevity

All experiments were performed under biosafety level 3 (BSL-3) conditions in the insectary at the Australian Animal Health Laboratory. Insectary conditions were maintained at 27.5°C and 70% in relative humidity with a 12hr light/dark cycle. ZIKV strain FSS13025 (Cambodia 2010, Genbank KU955593)^37^ or PRVABC59 (Puerto Rico 2015, GenBank KU501215) were used for viral challenge experiments. Both belong to the Asian/Pacific/American clade and were passaged once in C6/36 cells and twice in Vero cells before using for mosquito infections. WT (Higgs or Liverpool strain) and transgenic (confirmed by red fluorescence in the eye) mosquitoes were infected with ZIKV as previously described ^54^. Briefly, female mosquitoes were challenged with a chicken blood meal spiked with ZIKV (TCID_50_ 10^6^/mL) through chicken skin membrane feeding. Blood-fed female mosquitoes were sorted and maintained at standard conditions in an environmental cabinet with sugar *ad libitum*. For infection rate and virus titer, mosquito midguts were collected at 4 dpi. For dissemination and transmission rate, mosquito saliva, midguts, and carcasses were collected at 14 dpi. Mosquito saliva was used to determine viral titers using TCID_50_ assay on Vero cells. Midguts and carcasses were used to determine presence of viral RNA using RT-qPCR against ZIKV NS5 ^54^ (Supplementary Table 5). Mosquito viral challenge, processing, saliva testing, and molecular analyses of infection and dissemination were carried out as previously described ^54^. ZIKV infection rate was defined by the number of midguts (4 dpi) found positive for viral nucleic acid over tested midguts. Similarly, the dissemination rate was calculated by the number of carcasses (14 dpi) testing ZIKV positive by qPCR. Transmission rate was defined by the number of TCID_50_ positive saliva samples over the number tested. For each experiment, data from three replicates was pooled. The average TCID_50_ values were compared by two-tailed unpaired t test. To measure fitness after infection, blood-fed ZIKV-infected females were quickly sorted out after CO_2_ anaesthesia and housed in waxed cardboard cup 250 ml containers with a maximum of 25 mosquitoes. Mosquitoes were maintained at standard conditions for 14 days with 10% sugar solution *ad libitum*. Dead mosquitoes were counted daily. Females surviving at day 14 were marked as censored (status=0) in the database for survival analysis, which was performed using the GraphPad Prism software (GraphPad Software, La Jolla California, USA). The Mantel-Cox test was used to compare the survival of infected mosquitoes at 14 dpi.

### Generation of *w*Mel *Wolbachia* Line and Infection Assay

Eggs of *Ae. aegypti* infected with the *Wolbachia* strain *w*Mel were obtained from the World Mosquito Program (Prof. Scott O’Neill Monash University). Higgs mosquitoes infected with *w*Mel were generated by crossing *w*Mel+ females with males from the Higgs line, and the resulting offsprings were used for ZIKV infections experiments. At the end of the experiment, *Wolbachia* infection status of these mosquitoes was tested using PCR with primers specific for *w*Mel detection ^55^ (Supplementary Table 5). The PCRs indicated presence of *w*Mel in >90% of mosquitoes, and only results from these positive mosquitoes were used for further analysis.

### Mouse Transmission Assays

All experiments were performed under biosafety level 3 (BSL-3) conditions in the insectary at NHRI. Insectary conditions were maintained at 29°C and 80% relative humidity with a 12 hr light/dark cycle, and mosquitoes were maintained as previously described ^42^. For experimental assays, transgenic anti-ZIKV mosquitoes were outcrossed to WT (Higgs strain) for a generation to obtain heterozygotes. ZIKV strain PRVABC59 (Puerto Rico 2015, GenBank KU501215) was used for viral challenge experiments. It was obtained from the Taiwan Center for Disease Control, and maintained/amplified as previously described ^42^. For direct ZIKV infection, 7–10 day-old female mosquitoes were inoculated with 200 plaque forming units (pfu) of ZIKV by thoracic injection as previously described ^42^ and maintained under standard housing conditions for 7 days prior to their use in assays. Infection via artificial membrane blood feeding was carried out as described above, and infected mosquitoes were then maintained under standard conditions for 14 days prior to their use in transmission assays. Viral titers were measured at 7 dpi (for thoracic injection infections) or 14 dpi (for membrane blood feeding infection) by plaque assay as previously described ^42,56^. Briefly, 2×10^5^ cells/well of Vero cells (a kind gift from Dr. Guann-Yi Yu) were incubated for one day (in serum-free 1xDMEM medium (HyClone, SH30022), at 37°C) before being infected with ZIKV. At two hours post infection, unbound virus particles were removed, and cells were gently washed by PBS and overlaid with 3 ml of 1xDMEM medium containing 2% FBS (Gibco, 16000044), 10 mM HEPES, 10nM sodium pyruvate, 2mM L-Glutamine (Gibco, 25030081), 1x Penicillin-Streptomycin (Gibco, 15140122), and 1% Methyl cellulose (Sigma, M0512-250G). The infected cells were then incubated at 37°C and 5% CO_2_ for 4 days until plaque formation. Cells were fixed and stained with 0.5mL crystal violet/methanol mixed solution (ASK®Gram Stain Reagent) for 2 hours, and washed with H_2_O. Number of plaques was then calculated, and viral titers were determined as plaque forming units per mosquito and were compared by one-way ANOVA.

All mouse-related experiments were conducted in compliance with the guidelines of the Laboratory Animal Center of NHRI. The animal protocol (NHRI-IACUC-105111) was approved by the Institutional Animal Care and Use Committee of NHRI, according to the Guide for the Care and Use of Laboratory Animals (NRC 2011). Management of animal experiments and animal care and use practices of NHRI have been accredited by the AAALAC International. *Stat1* -/- (C57BL/6 background) mice were provided by Dr. Guann-Yi Yu (NTU, Taiwan). Both male and female mice between the ages of 11-12 weeks were used in the study.

Mosquito-mediated ZIKV mouse infections were carried out as previously described ^42,56^. Briefly, mice were anesthetized with Ketalar (100 mg/Kg, Pfizer, New York, NY) via intraperitoneal injection, and their ventral surfaces were shaved. Then, mice were placed on top of a polyester mesh covering a mosquito-housing cage that permitted female mosquitoes to take a blood meal. Female mosquitoes were starved for 10h before they were allowed to take blood meals from mice, and each mouse was fed on by 6–11 mosquitoes. Mouse body weight and mortality were then recorded for 6-30 days. Mouse weights were compared by the Mann Whitney test to evaluate for significant weight loss.

### Fitness Assessment and Conditions

To determine if the anti-ZIKV transgene confers a fitness cost, several fitness parameters were evaluated in WT and transgenic mosquitoes. Evaluation of all experimental and control replicates were performed simultaneously. Insectary conditions were maintained at 28°C and 70-80% in relative humidity with a 12hr light/dark cycle. To assess larval to pupal development time, eggs were vacuum hatched and larvae were distributed into pans (50 larvae per pan) containing 2.5L of ddH_2_O and 0.6mL of fish food slurry. To determine the development time of transgenic and WT control mosquitoes, 4th instar larvae were sorted according to fluorescence phenotype and reared until pupation. Pupae were collected and counted every day until no pupae were left. To assess female fertility and fecundity, 90 females were mated to 20 WT males in a cage. After four days, females were blood fed and individually transferred into plastic vials filled with water and lined with egg paper. After three days, egg papers were collected, and eggs were counted and vacuum hatched in 9-ounce plastic cups. Starting on the fourth day, larvae were counted every day until no larvae were present. Female fecundity refers to the number of eggs laid per female, and fertility reflects the number of eggs hatching to produce larvae. To measure male mating success, fecundity, and fertility, one transgenic or control male was mated to five WT females in a single cup filled with water and lined with egg paper. Three days post blood meal, cups were checked for the presence of eggs, which were collected, counted, and hatched. Hatched larvae were then counted every day until no larvae were present. Male mating success was calculated as the percentage of single male outcrosses that produced larvae. Fecundity was measured as the number of eggs laid per cup; fertility was determined by the number of hatching larvae in each cup. To asses wing length as a proxy for body size, images of mosquito wings were taken with a Leica M165 FC microscope (Leica Microsystems). Wing length measurements were done by using the measurement tool on the Leica Application Suite X, measuring from the axial incision to the intersection of the R 4+5 margin. Finally, to assess mosquito longevity, equal numbers of male and female experimental or WT mosquitoes were placed in medium sized cages (in triplicate). Mosquitoes that died were counted and removed daily until all mosquitoes had died. Statistical analyses were performed using the GraphPad Prism software (GraphPad Software, La Jolla California, USA). Means were compared using unpaired t tests with Welch’s correction except for male mating success (no Welch’s correction). Analyses of mosquito survivorship utilized the Mantel-Cox test. *P*-values>0.05 were considered non-significant.

### Confirmation of Transgene Zygosity

To molecularly confirm zygosity of transgenic mosquitoes, mosquito heads were homogenised using bead-beater for DNA extraction in 30 ul extraction buffer (1x Tris-EDTA, 0.1M EDTA, 1M NaCl and 2.5 uM proteinase K), and incubated at 56°C for 5 minutes and then at 98°C for 5 minutes. PCR was then performed on each line to detect the presence of the transgene by pairing a *piggyBac* primer with a genomic primer as follows: primers 1018.S46 and 991.5R2 for TZIK-A, 1018.S26 and 991.3F2 for TZIK-B, 1018.S8 and 991.5R1 for TZIK-C, and 1018.S50 and 991.3F2 for TZIK-D (Supplementary Table 5). To determine zygosity, we amplified the WT locus of each transgenic line using corresponding forward and reverse primers listed in Supplementary Table 5. A PCR kit (ThermoFisher Scientific #F553S) with 57°C annealing temperature and standard protocols was used for all PCRs.

### Data Availability Statement

All sequencing data associated with this study are available from NCBI sequence read archive (SRA) accession ID: SRP150144; BioProject ID: PRJNA475410. Complete annotated plasmid sequence and DNA is publically available at Addgene (plasmid #104968). Transgenic mosquitoes will be made available by corresponding author upon request.

## Supplemental Figures and Tables

**Supplementary Figure S1.**
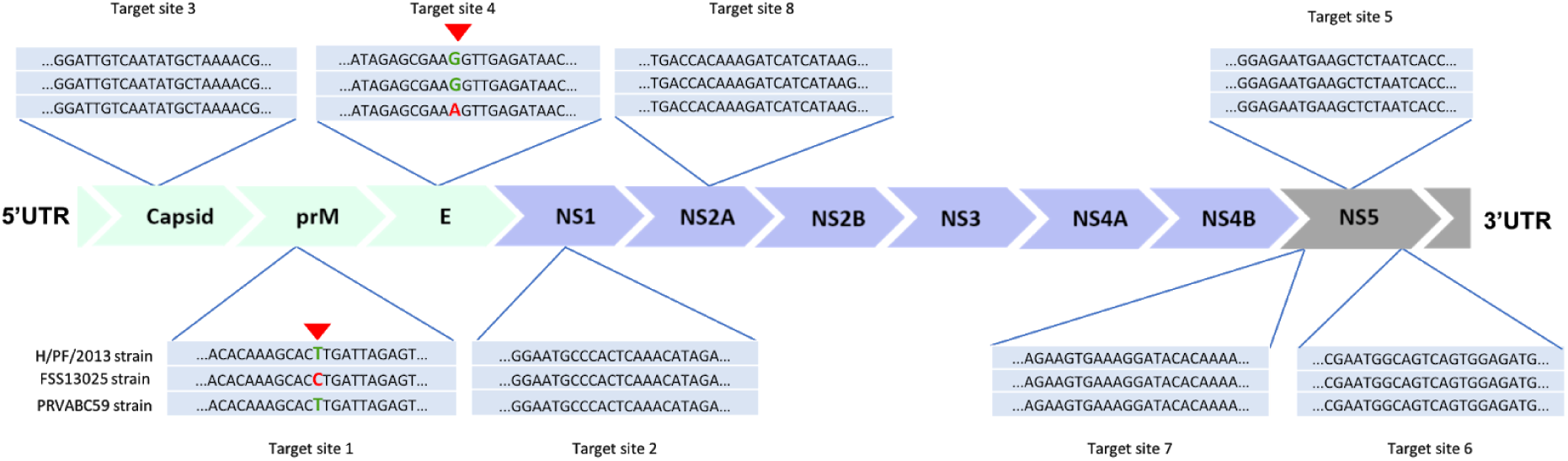
miRNA target site conservation between ZIKV strains H/PF/2013, FSS13025, and PRVABC59. miRNA target sites between the ZIKV strain used for miRNA target selection (H/PF/2013, top sequence) and the strains used for mosquito challenges (FSS13025, middle sequence; PRVABC59, bottom sequence) are highly conserved, with only one base pair mismatch in one target site in each strain (shown in red).

**Supplementary Figure S2:**
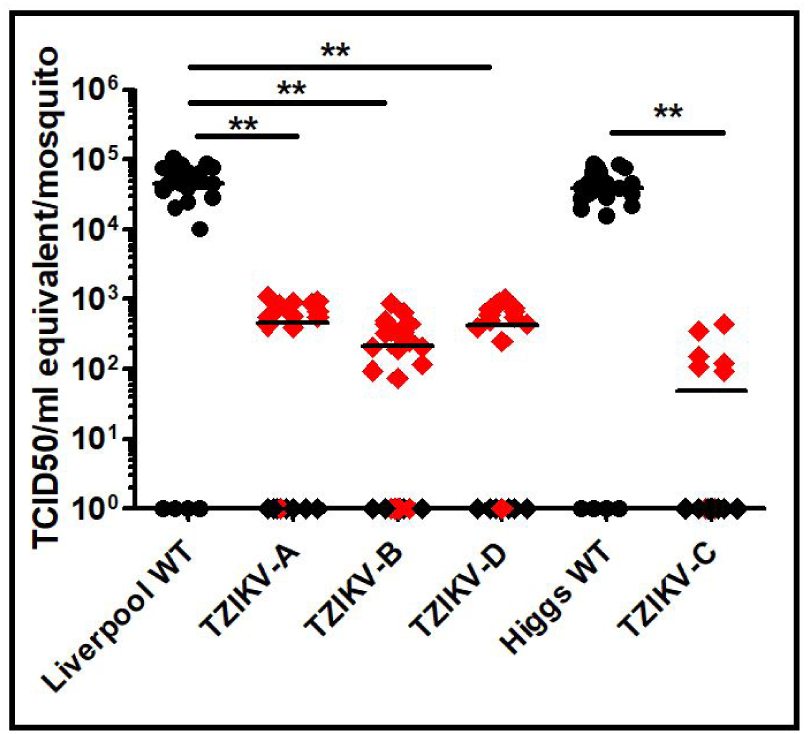
Effect of anti-ZIKV transgene on ZIKV titres in four independent mosquito lines. ZIKV virus titres in wildtype (Liverpool WT and Higgs WT), anti-ZIKV transgenic mosquito lines (TZIKV-A, TZIKV-B, TZIKV-C, TZIKV-D) following a blood meal infected with a Cambodian (FSS13025) are shown. Virus titres (TCID50/ml equivalent) from mosquito midgut (day 4 post infection) of WT and transgenic mosquitoes were determined using real-time RT-qPCR and calculated using previously published methods. Circles represent WT mosquitoes; black diamonds represent anti-ZIKV Hm transgenic mosquitoes; red colored diamonds represent anti-ZIKV Ht transgenic mosquitoes. Horizontal bars represent the mean virus titer. Mantel-Cox test was used for statistical analysis. **represents p<0.001.

**Supplementary Figure S3.**
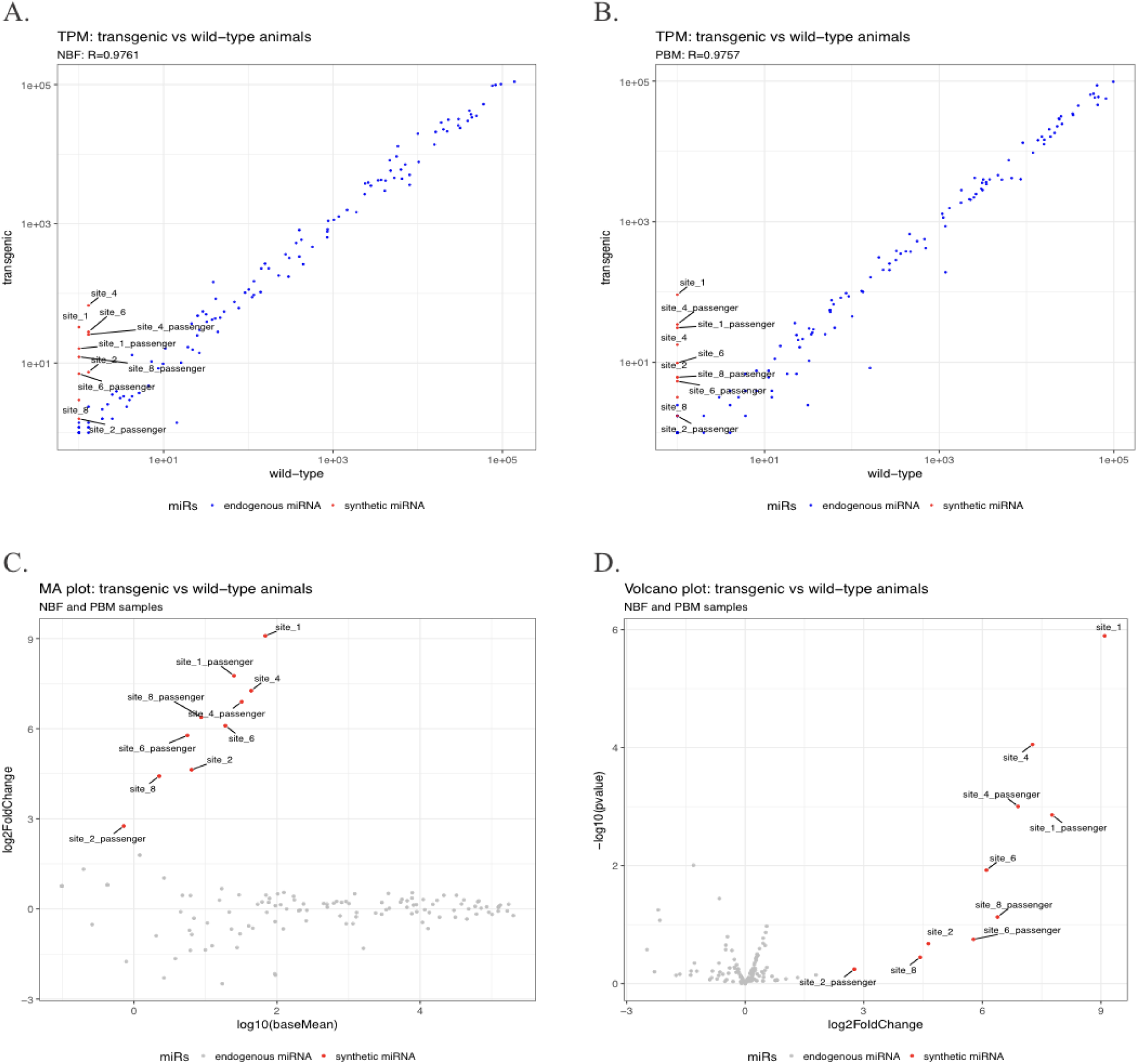
Differential expression analysis of miRNAs from WT and TZIKV-C mosquito midguts. TPM (transcripts per million) values for transgenic versus WT animals without a blood meal (**A**) and 24 hours after a blood meal (**B**) are shown. Expression of synthetic miRNAs does not affect expression levels of endogenous miRNAs significantly (correlation coefficients of 0.9761 and 0.9757, respectively). MA (log2FoldChange vs. baseMean)(**C**) and ‘volcano’ (-log10 (p-value) vs. log2FoldChange))(**D**) plots demonstrate that only synthetic miRNAs are differentially expressed between WT and transgenic animals.

**Supplementary Figure S4.**
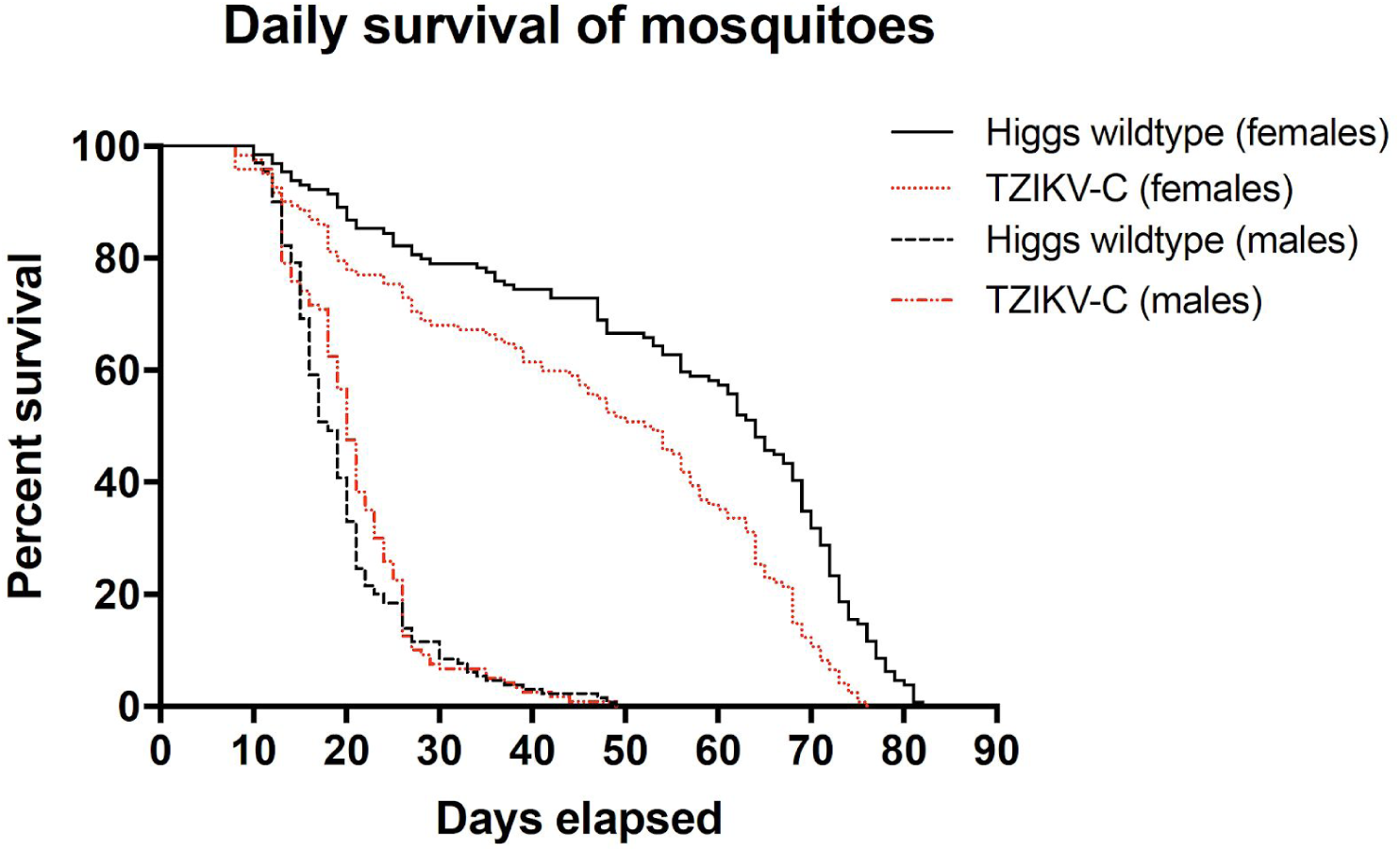
Survivorship curve of WT and TZIKV-C male and female mosquitoes. The x-axis indicates the number of elapsed days after the start of the experiment, and the y-axis indicates the percent of mosquitoes surviving on each elapsed day. Each line represents accumulated results from 120-130 adult mosquitoes combined from 3 biological replicates.

**Supplementary Table S1.** Quantification of endogenous and engineered miRNA expression on read and UMI (Unique Molecular Identifiers) levels in WT and TZIKV-C mosquitoes prior to blood meal (NBF) and 24 hr post blood feeding (PBM). Both raw read or UMI counts and normalized TPM (Transcripts Per Million) values are shown.

**Supplementary Table S2.** The survivorship of ZIKV-infected TZIKV-C mosquitoes at 14 days post infection (dpi). WT, *w*Mel+, and TZIKV-C mosquitoes infected with ZIKV strain FSS13025 or PRVABC59 were assessed for survival at 14 dpi. The mean percentage±SEM of surviving mosquitoes and number of mosquitoes tested (in parentheses) are reported. No assay was performed for *w*Mel mosquitoes infected with strain FSS13025. The Mantel-Cox test was used to compare the survival of infected WT, *w*Mel (for PRVABC59 strain only), and anti-ZIKV mosquitoes.

| Virus strain | Mosquito strains |  |  | P-value |
| --- | --- | --- | --- | --- |
|  | WT | WT<br>*w*Mel | TZIKV-C |  |
| FSS13025<br>(Cambodia) | 64.1% ± 6.6 (53) | -- | 73.6% ± 6.1 (53) | 0.2528 |
| PRVABC59<br>(Puerto Rico) | 92.1% ± 3.0 (76) | 86.9% ± 5.0 (46) | 77.1% ± 7.0 (35) | 0.0817 |

**Supplementary Table S3.**
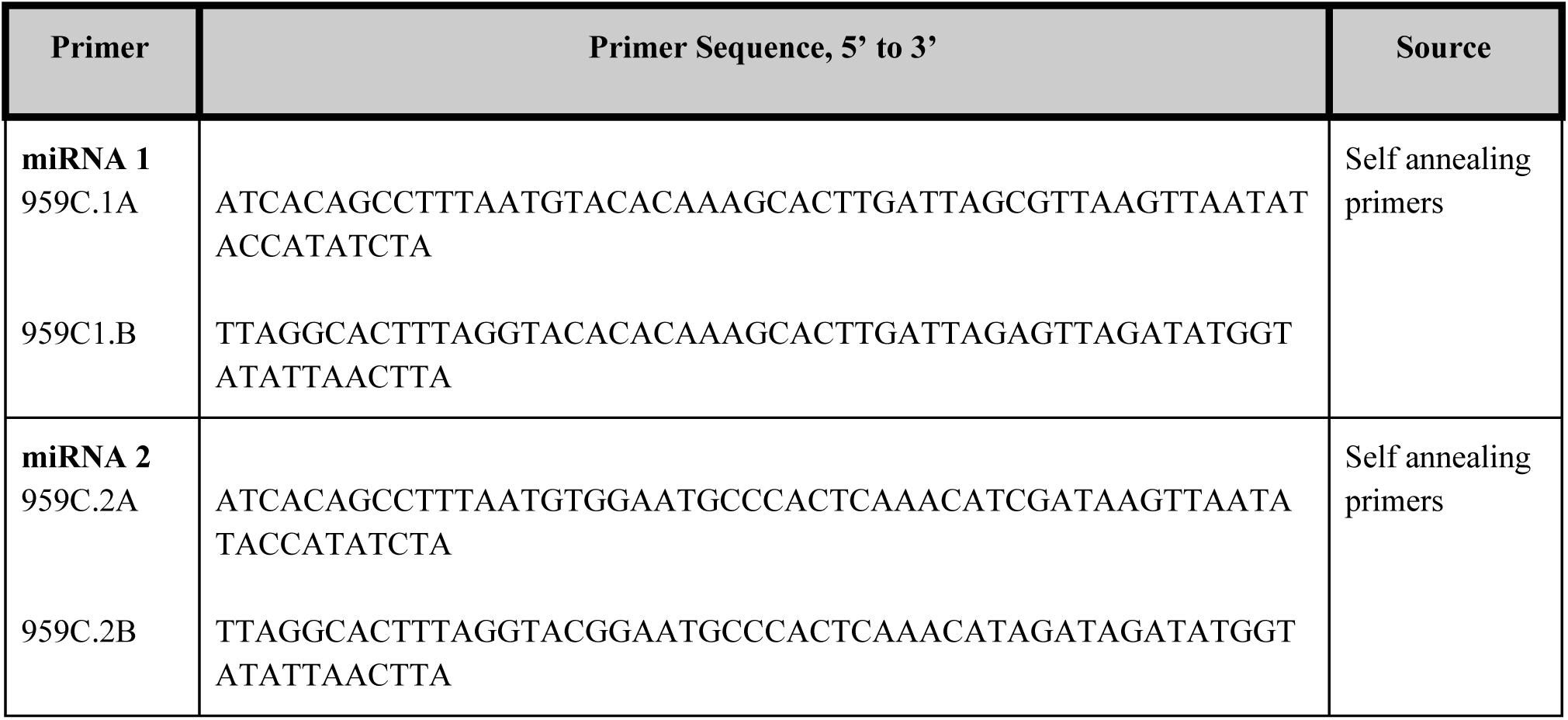
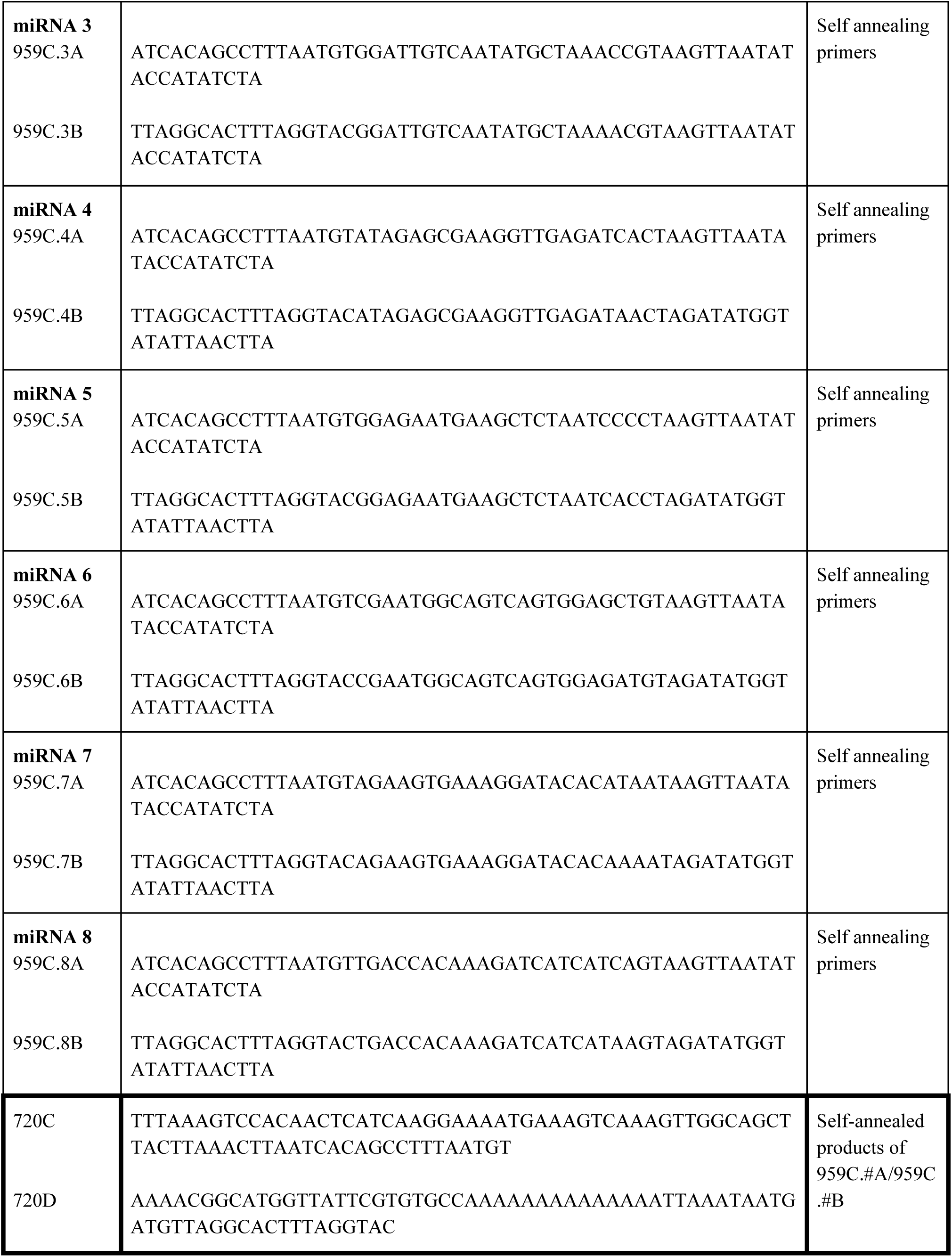
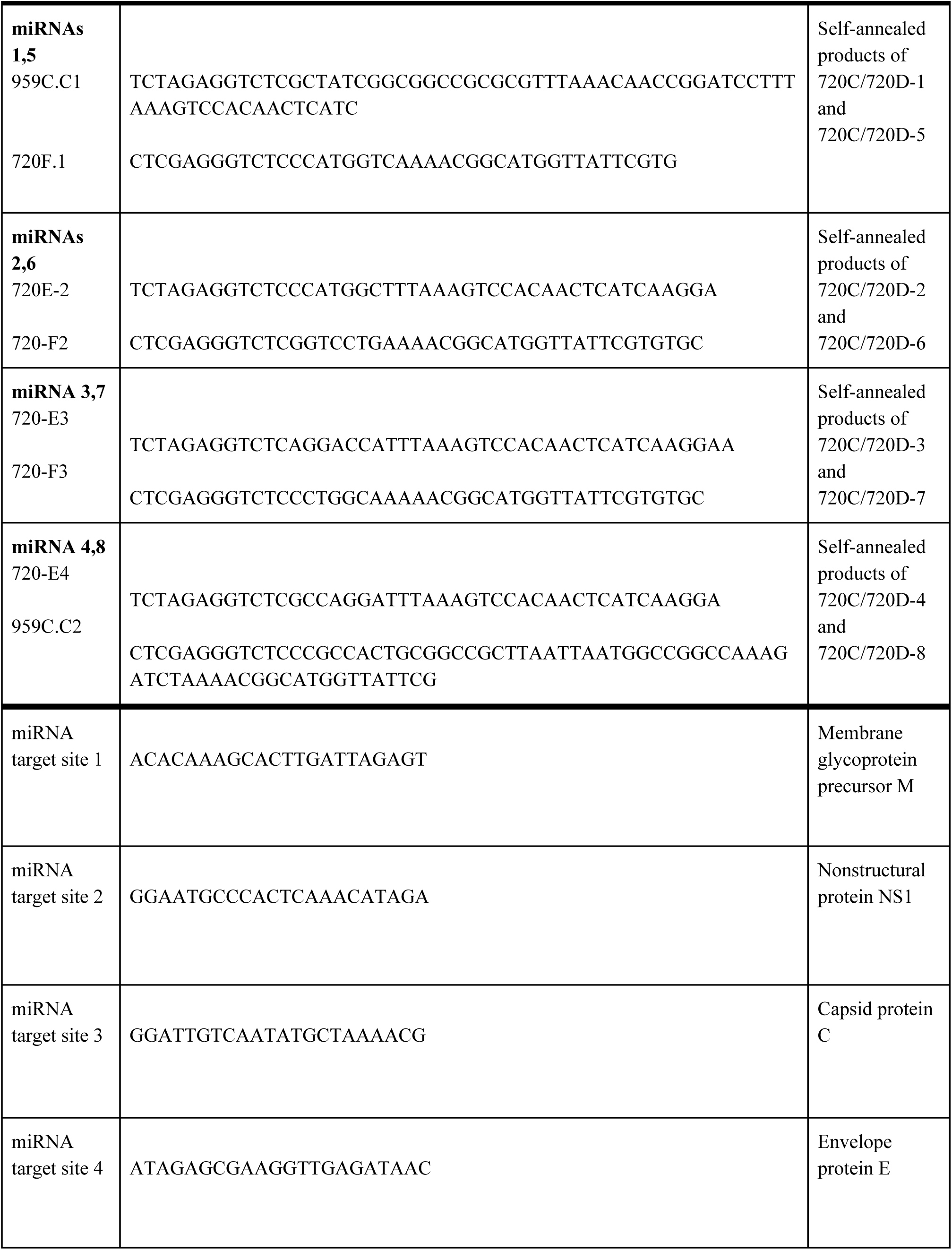
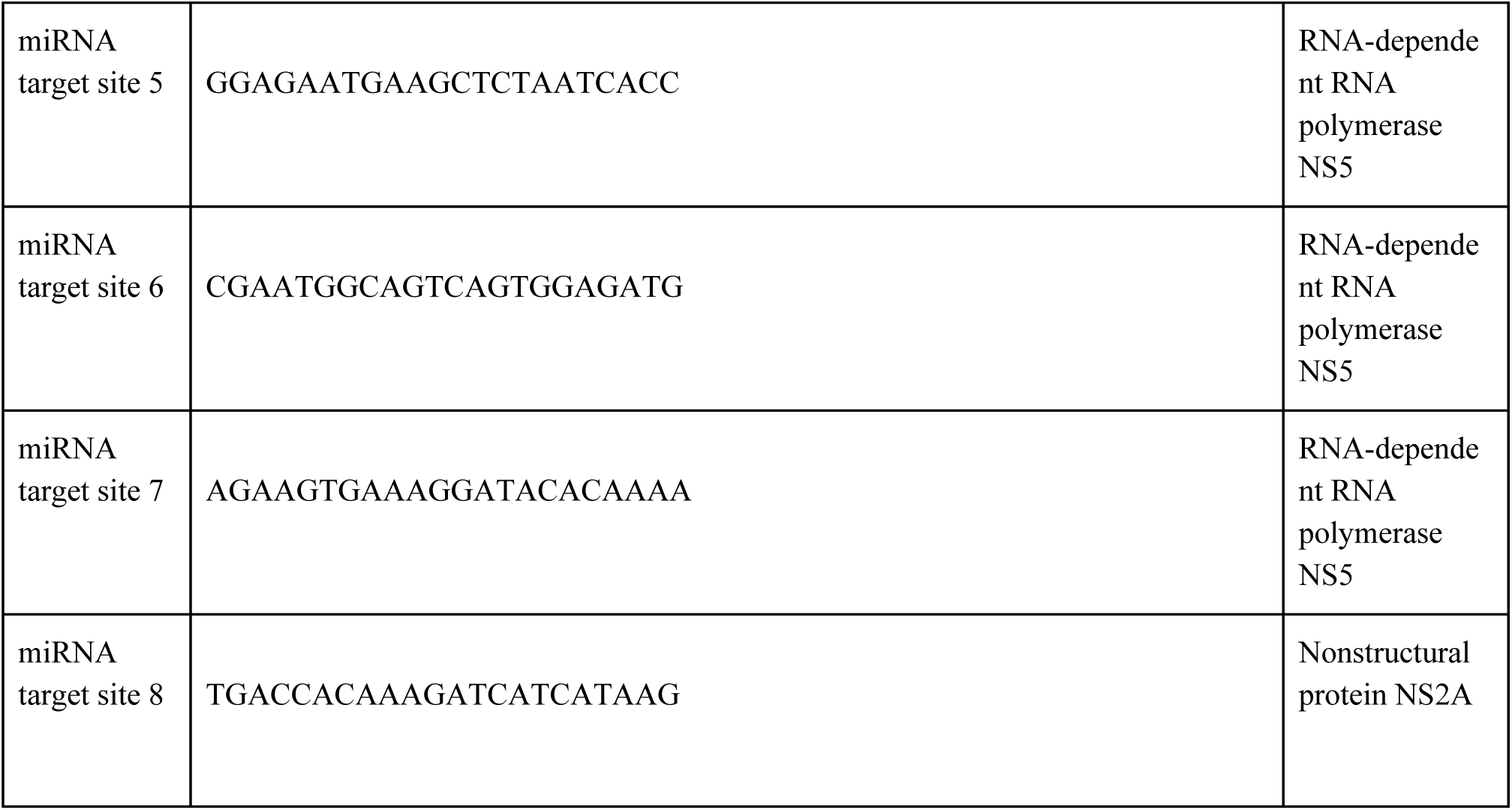
Primer sequences and miRNA target sites utilized to generate synthetic miRNA constructs used in this study. Self annealing primers are listed first, and consist of forward and reverse target site sequences flanking the stem loop region of the synthetic miRNA. Primers amplifying flanking regions, BsaI cut sites, and multiple cloning site are listed below.

**Supplementary Table S4.**
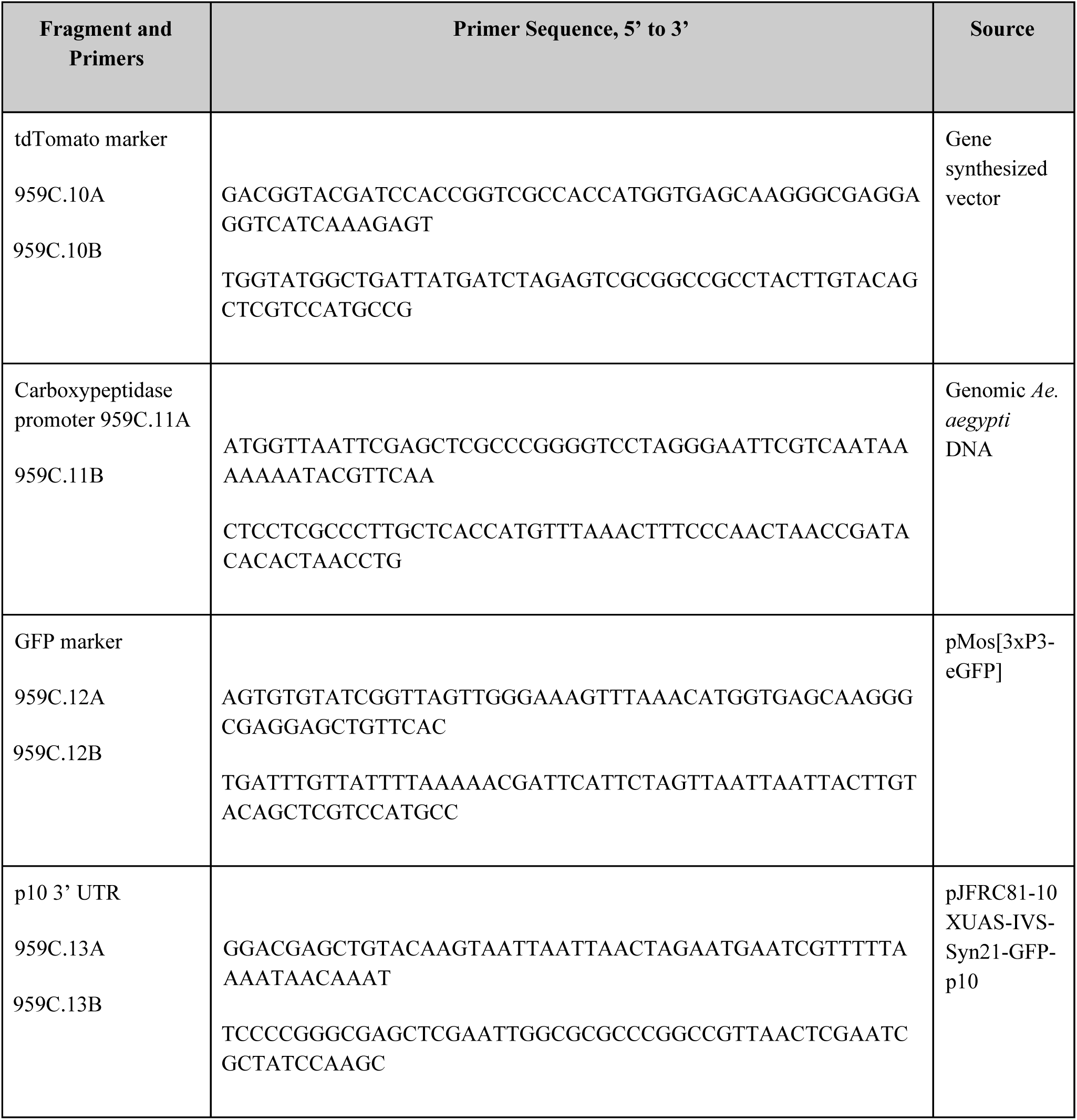
Primers used to assemble plasmid OA959C (the anti-ZIKV transgene).

**Supplementary Table S5.**
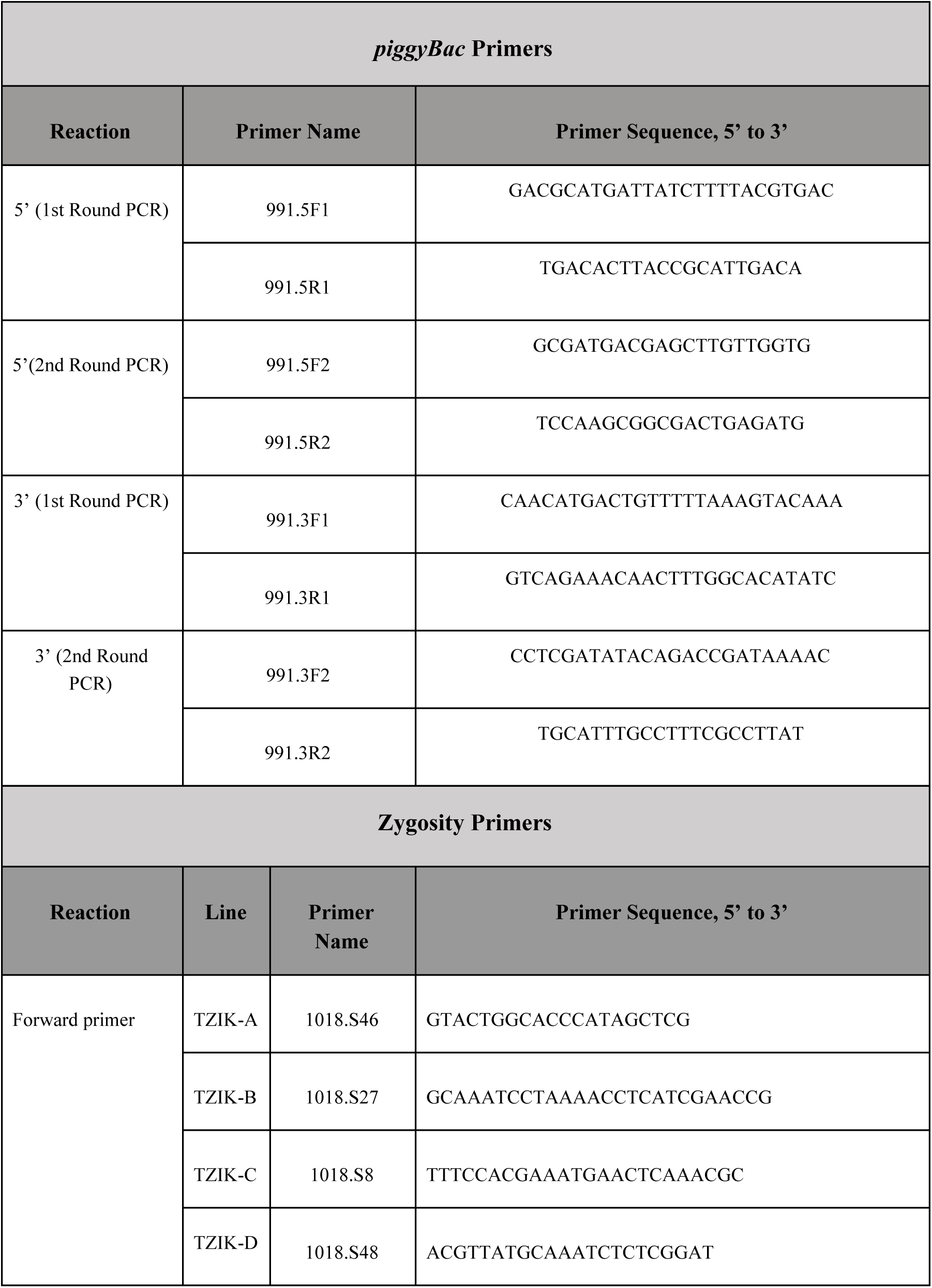
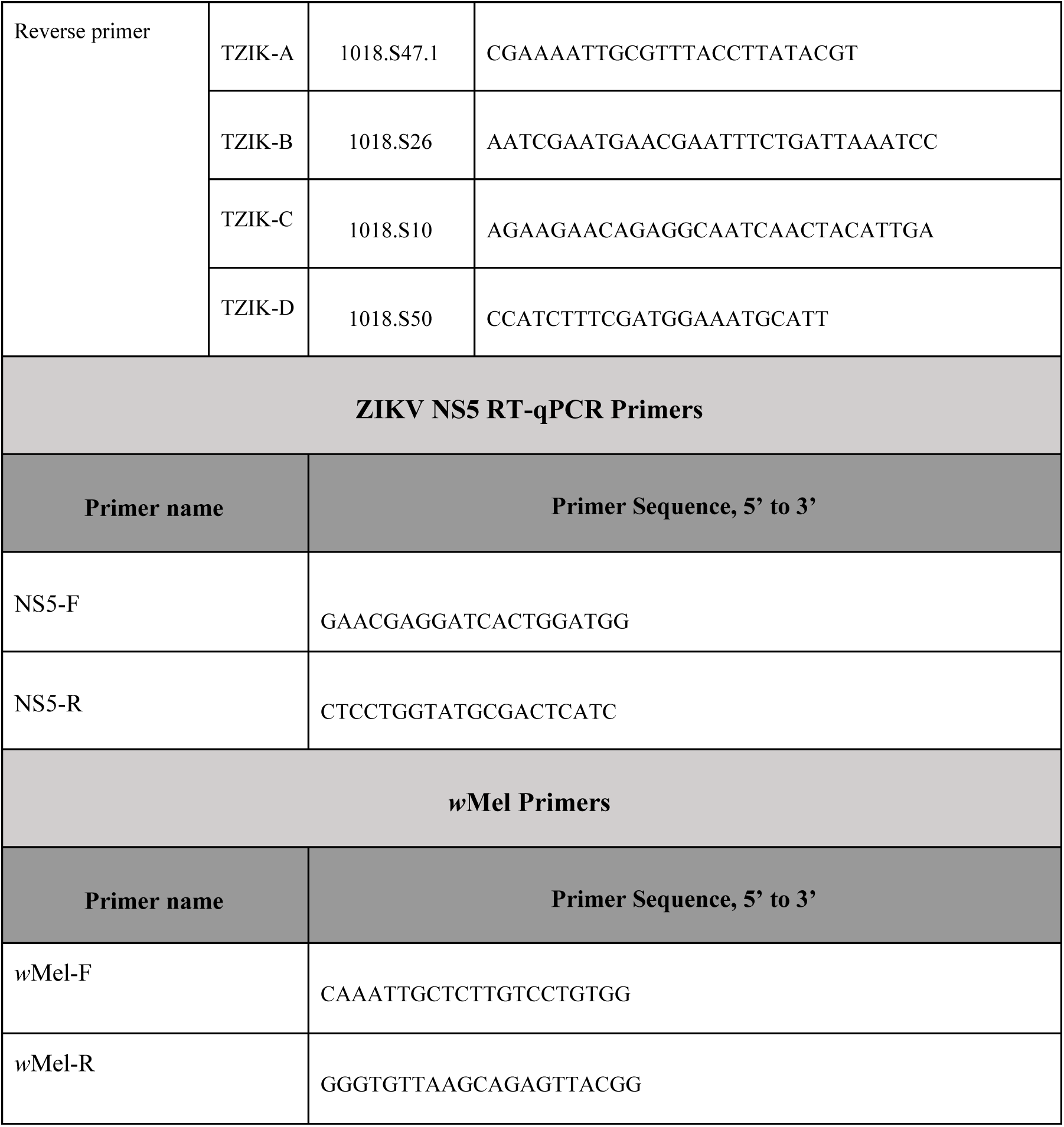
Diagnostic primers used for inverse PCR (iPCR) assays, zygosity confirmation, ZIKV NS5 RT-qPCR, and *w*Mel infection confirmation.

